# CEBPA restricts alveolar type 2 cell plasticity during development and injury-repair

**DOI:** 10.1101/2023.10.10.561625

**Authors:** Dalia Hassan, Jichao Chen

**Affiliations:** Department of Pulmonary Medicine, the University of Texas MD Anderson Cancer Center, Houston, Texas 77030, USA; The University of Texas MD Anderson Cancer Center UTHealth Graduate School of Biomedical Sciences, Houston, Texas 77030, USA

**Keywords:** cell plasticity, transcriptional control, cell fate, epigenome, lung development and regeneration, single-cell multiome

## Abstract

Cell plasticity theoretically extends to all possible cell types, but naturally decreases as cells differentiate, whereas injury-repair re-engages the developmental plasticity. Here we show that the lung alveolar type 2 (AT2)-specific transcription factor (TF), CEBPA, restricts AT2 cell plasticity in the mouse lung. AT2 cells undergo transcriptional and epigenetic maturation postnatally. Without CEBPA, both neonatal and mature AT2 cells reduce the AT2 program, but only the former reactivate the SOX9 progenitor program. Sendai virus infection bestows mature AT2 cells with neonatal plasticity where *Cebpa* mutant, but not wild type, AT2 cells express SOX9, as well as more readily proliferate and form KRT8/CLDN4+ transitional cells. CEBPA promotes the AT2 program by recruiting the lung lineage TF NKX2-1. The temporal change in CEBPA-dependent plasticity reflects AT2 cell developmental history. The ontogeny of AT2 cell plasticity and its transcriptional and epigenetic mechanisms have implications in lung regeneration and cancer.

## INTRODUCTION

Cell plasticity, semantically defined as a cell’s ability to become a different cell type, determines the resilience and reaction of cells to genetic and environmental perturbations. Theoretically, nearly every cell has the same DNA blueprint to become any other cell type, as demonstrated by reprogramming of fibroblasts into pluripotent stem cells^1^. In tissues, more plasticity is desirable for progenitor/stem cells to fuel their more differentiated lineages and for direct cell fate switch during trans-differentiation, but can be hijacked during tumorigenesis^2^. Less plasticity is associated with developmental differentiation toward specialized physiology, as well as with ageing^3^.

During development, progenitors gradually confine themselves first to particular germ layers, then anteroposterior dorsoventral locations and organs, and finally cell types within, gaining cell-type specificity and losing plasticity – apparently two sides of the same coin. Analogous to evolutionary speciation, this developmental history sets the time of divergence among cell types and the rates afterwards – trajectories potentially predictive of the direction and extent of cell plasticity. The graduality of development is punctuated by points of no return, beyond which cells cannot revert to their developmental ancestors, as illustrated by closure of the neonatal regenerative window for the mammalian heart^4^. These points of no return often differ from the points of cell fate specification, highlighting the importance of post-specification maturation and the discordance between the said two sides of the specificity-plasticity coin. Therefore, factors traditionally studied as cell-fate promoting need to be separately evaluated for plasticity restricting. Such studies of reactivating developmental plasticity and overcoming points of no return can inform tissue regeneration and are medically important.

The definitional dependence of cell plasticity on cell types results in operational difficulties, as the spectrum of variable states within a cell type often bleeds into related cell types, especially during development and upon injury, introducing uncertainty in cell types and thus plasticity. However, the continuum of cell types can be objectively quantified by molecular profiling of the transcriptome, epigenome, proteome, etc., especially when done on a single-cell level. Accordingly, this study defines cell plasticity as gains in molecular features characteristic of other cell types.

Building on our prior transcriptional and epigenetic studies of the mouse lung epithelium, this study focuses on the alveolar type 2 (AT2) cells that originate from SOX9 embryonic progenitors and become facultative stem cells, which secret pulmonary surfactants at baseline, but self-renew and give rise to the gas-exchanging alveolar type 1 (AT1) cells upon injury^5,6^. This life cycle of AT2 cells provides an experimental paradigm to probe the ontogeny of cell plasticity and associated transcriptional regulators. We show that AT2 cells use a cell-type-specific transcription factor (TF) CEBPA to restrict their plasticity at neonatal and mature stages, but reactivate the developmental plasticity upon respiratory virus infection. Mechanistically, CEBPA recruits the lung lineage TF NKX2-1 to promote the AT2 cell program and indirectly represses the SOX9 progenitor program, within the confinement of their developmental history.

## RESULTS

### Postnatal transcriptomic and epigenomic maturation of AT2 cells is separate from their embryonic specification

To explore how cell plasticity can be shaped by development, we first delineated the associated molecular progression of AT2 cells using scRNA-seq and scATAC-seq. Mouse lung epithelial cells profiled by scRNA-seq at 12-time points spanning from embryonic day (E) 14.5 to 15-week adult stages, clustered by time and cell types (Fig. 1A). AT2 cells clustered separately from AT1 cells only by E18.5, indicating measurable specification from their E14.5 and E16.5 SOX9 progenitors (Fig. 1A). Nascent AT2 cells at E18.5 were distinct from their neonatal and adult counterparts, suggesting maturation following specification, which was supported by Monocle trajectory analysis (Fig. 1B). Eighty-two genes eventually specific to AT2 cells were designated as early versus late AT2 genes based on expression, however low, in >25% of E16.5 SOX9 progenitors^7^ (Fig. 1C). By comparison, 10 out of 41 late AT2 genes, including immune response genes (*Lyz2*, *Lrg1*, *Chil1*, and *H2* paralogs), only reached maximal expression (>4-fold increase from E18.5 to 15-week) during postnatal maturation (Fig. 1C).

**Figure 1.**
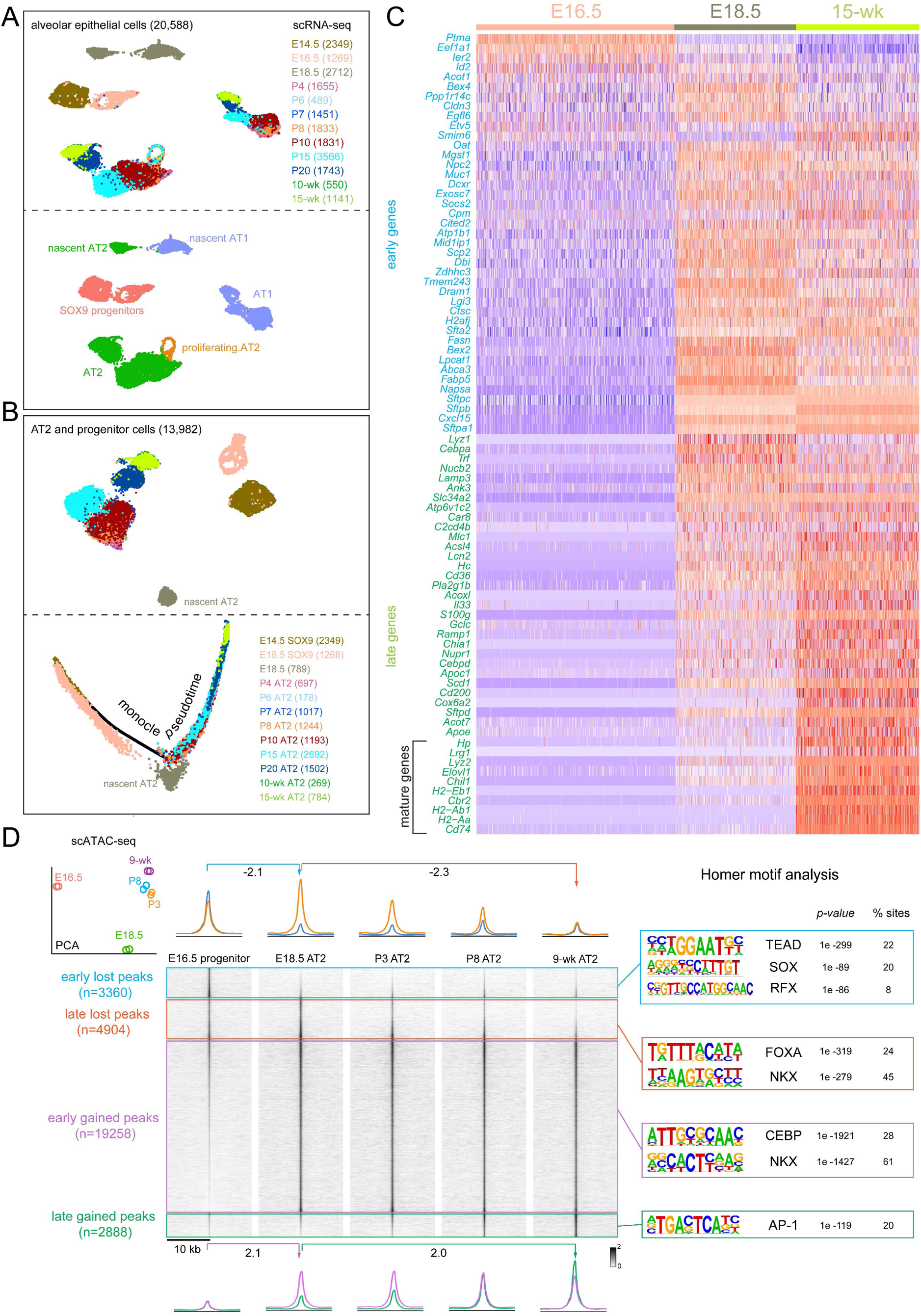
AT2 cells undergo transcriptomic and epigenomic maturation postnatally, separate from their embryonic specification. (**A**) Aggregated scRNA-seq UMAPs of alveolar epithelial cells of wild type lungs from 12 time points color coded by time (top) or cell type (bottom). Cell numbers are in parenthesis. (**B**) Top: UMAP of AT2 and SOX9 progenitor cells subset from in (**A**), colored coded by time, and their monocle pseudotime analysis (bottom) showing molecular progression from E14.5 SOX9 progenitors through E18.5 nascent AT2 to 15-wk mature AT2 cells. Cell numbers are in parenthesis. (**C**) Expression heatmap of 82 AT2-specific genes, classified as early if present in at least 25% of cells at E16.5. The remaining late genes are considered mature if the fold increase from E18.5 to 15-wk is more than 4. (**D**) Principal component analysis (PCA) of scATAC-seq pseudobulk duplicates showing the distinct epigenome of E18.5 nascent AT2 cells. ScATAC-seq heatmaps and profile plots categorized and color coded as early lost (E16.5 vs E18.5), late lost (E18.5 vs 9-wk), early gained (E16.5 vs E18.5) and late gained (E18.5 vs 9-wk) and their Homer motif analysis. Peak numbers are in parenthesis. See Table S1 for raw data.

Supporting this transcriptional post-specification maturation, scATAC-seq sampling of 5 developmental time points from E16.5 to 9-week adult revealed epigenomic maturation of AT2 cells that occurred after their E18.5 specification (PCA plot in Fig. 1D). Accordingly, we classified differential ATAC peaks into lost and gained groups, each with early and late subgroups separated at the E18.5 time point of specification (Fig. 1D). Although ATAC peaks often correlate with RNA transcripts and may act over long-range and on high-order chromatin structure, they measure individual regulatory regions without averaging over the whole gene, predict regulatory TF motifs, and bypass the confounding issue of perdurance in RNA-seq. As shown in Fig. 1D, the early lost peaks coincided with AT2 specification at E18.5, were near progenitor and AT1 genes (e.g. *Adamts18* and *Pdpn*, respectively), and contained SOX and TEAD motifs, likely reflecting the termination of the SOX9-mediated progenitor program and the low level of YAP/TAZ/TEAD-mediated AT1 program in E16.5 progenitors^7,8^. The late lost peaks decreased postnatally, were near stem cell genes (e.g. *Klf4*, *Id4*, and *Etv6*)^1,9,10^, and contained FOXA and NKX motifs, correlating with the postnatal decrease in AT2 cell proliferation and potential progenitor-specific functions of FOXA2 and NKX2-1. Conversely, the majority of differential peaks (19,258 out of 30,410; 63%) were in the early gained subgroup and were near AT2 genes and enriched for CEBP and NKX motifs, which were examined in detail in this study. Last, the late gained peaks followed the RNA kinetics of AT2 cell maturation (Fig. 1C), were near the corresponding genes (e.g. *H2-Aa* and *Cd74*), and contained the AP-1 motif. Supporting the epigenomic distinction between specification and maturation, each of the 4 groups was associated with distinct biological pathways and unsupervised clustering of top 100 variable motifs predicted from scATAC-seq revealed lost, early gained, and late gained groups, dominated by SOX and TEAD, CEBP, and AP-1 motifs, respectively (Fig. S1A, S1B).

We reasoned that, while the gain in chromatin accessibility reflected AT2 cell differentiation, the concurrent loss in accessibility not only indicated the conclusion of prior cell fates, but also predicted available fates when AT2 cells became more plastic. Accordingly, TFs promoting AT2 cell differentiation might also restrict AT2 cell plasticity. Of particular interest was CEBPA, whose motif was enriched among the early gained peaks, as well as our published AT2-specific NKX2-1 ChIP-seq peaks^8^ (Fig. 1D). Compared to other CEBP family members, *Cebpa* was specific to AT2 cells and reached maximal expression coinciding with AT2 specification (Fig. S1C, S1D). Supporting this, CEBPA was absent on the protein level at E14.5 or E16.5 when branch tips consisted of SOX9+ progenitors, but was expressed from E18.5 on in a subset of epithelial cells that were cuboidal, SOX9-HOPX-LAMP3+, and thus nascent AT2 cells (Fig. 2A, S1E, S1F). Taken together, the time-course transcriptomic and epigenomic analyses of AT2 cells provide a molecular roadmap of their sequential specification and maturation and implicate CEBPA in AT2 cell differentiation and plasticity.

**Figure 2.**
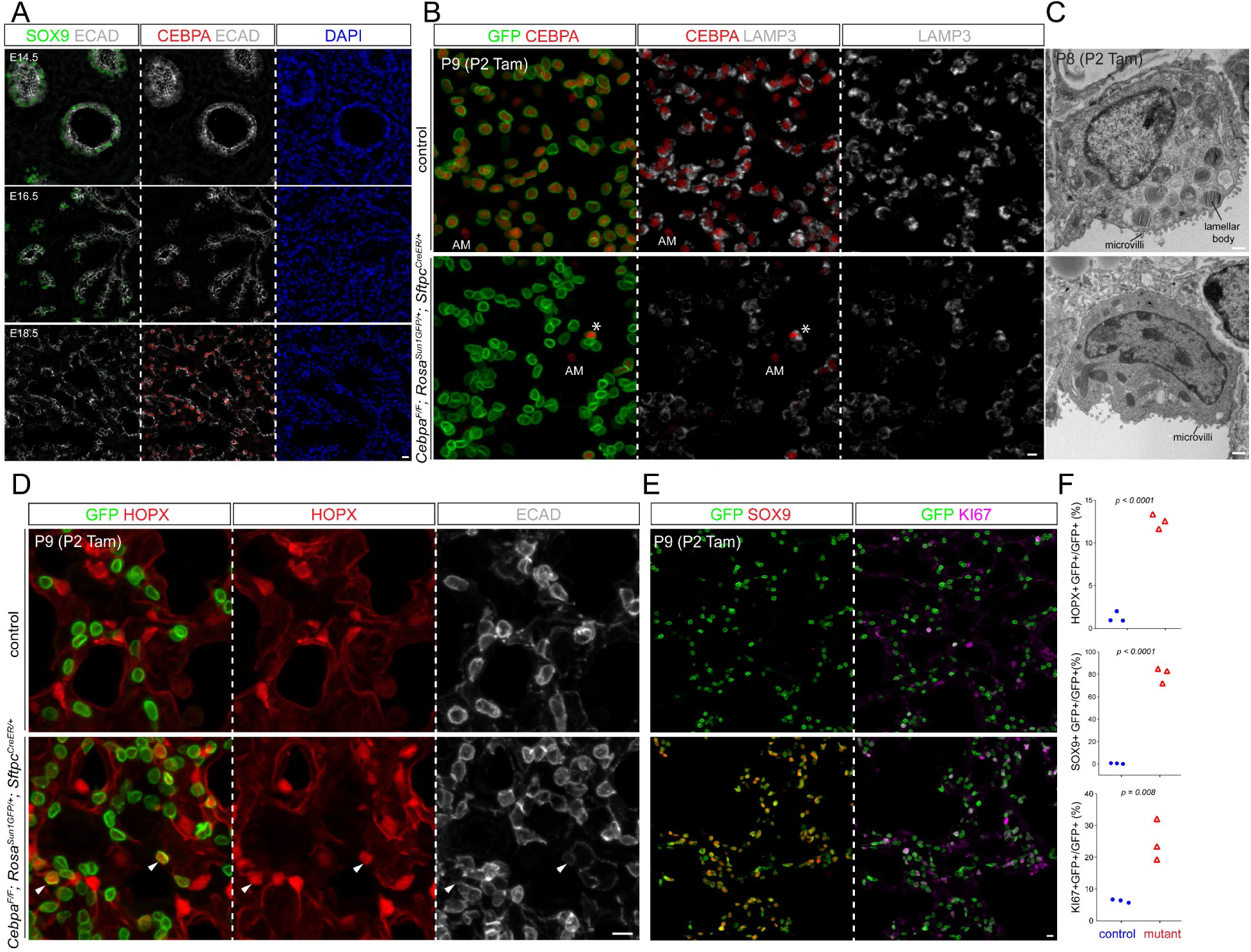
CEBPA promotes AT2 and suppresses progenitor programs in neonatal AT2 cells. (**A**) Confocal images of immunostained wild type lungs showing little CEBPA expression at E14.5 and E16.5 when branch tips are dominated by SOX9 progenitors but CEBPA expression in cuboidal cells outlined with E-Cadherin (ECAD) at E18.5. (**B**) Confocal images of immunostained neonatal AT2-specific *Cebpa* mutant and littermate control lungs showing loss of CEBPA in GFP+ recombined cells (asterisk: escaper), without affecting its expression in alveolar macrophages in the airspace (AM), and reduced LAMP3. Tam, 250 ug tamoxifen. Images are representative for at least three lungs (same for subsequent immunostainings). (**C**) Transmission electron microscopy (TEM) images showing reduction in lamellar bodies in mutant AT2 cells without affecting their apical microvilli. Tam, 250 ug tamoxifen. See Fig. S2B for more examples and quantification. (**D**) Confocal images showing lineage labeled mutant AT2 cells expressing an AT1 marker HOPX and no longer cuboidal (ECAD outline) (arrowhead). (**E**) Confocal images showing lineage labeled mutant AT2 cells ectopically expressing a progenitor marker SOX9 and a proliferation marker KI67. (**F**) Quantification of (**D**) and (**E**). Each symbol represents one mouse from littermate pairs (Student’s t-test). Scale: 10 um for all except for (**C**) 1 um. See Table S2 for raw data.

### CEBPA promotes AT2 and suppresses progenitor programs in neonatal AT2 cells

Although CEBPA had been studied in embryonic lungs^11^, to dissect the role of CEBPA throughout AT2 cell development, we generated an inducible AT2-specific knockout model *Cebpa^F/F^; Sftpc^CreER/+^; Rosa^Sun1GFP/+^*. To target AT2 cells shortly after specification, we induced Cre-recombination at a neonatal stage (P2) with an efficiency of 99% and specificity of 96% (1310 GFP+ cells from 3 mice) and deleted CEBPA in AT2 cells with an efficiency of 88% (1951 GFP+ cells from 3 mice), without affecting its normal expression in alveolar macrophages (Fig. 2B). By P9, *Cebpa* mutant AT2 cells had a drastic decrease in LAMP3 and a loss of IL33, both AT2 markers, compared to adjacent escapers of Cre-recombination or AT2 cells in the littermate control (Fig. 2B, S2A). Transmission electron microscopy showed that cuboidal/columnar epithelial cells in the *Cebpa* mutant lung often lacked lamellar bodies, a defining feature of AT2 cells, but still had characteristic apical microvilli (Fig. 2C, S2B).

Our previous study of the CEBPA equivalent in AT1 cells, YAP/TAZ/TEAD, showed activation of the alternative alveolar program^8^ and thus prompted us to examine the AT1 program in *Cebpa* mutant AT2 cells. To our surprise, only a small fraction of mutant AT2 cells expressed an AT1 marker HOPX (12.5% of 2049 GFP+ cells from 3 mice), lost LAMP3, and were no longer cuboidal as outlined by E-Cadherin, compared to a baseline of 1.2% in the control (1353 GFP+ cells from 3 mice) that possibly resulted from driver non-specificity/leakiness and normal neonatal conversion of AT2 to AT1 cells (Fig. 2D). Instead, we noticed adjoining *Cebpa* mutant cells reminiscent of SOX9 progenitors at embryonic branch tips (Fig. S2C). Remarkably, 80% of GFP+ cells in the mutant lung (1202 GFP+ cells from 3 mice) expressed SOX9, compared to 0.7% in the control (1370 GFP+ cells from 3 mice) (Fig. 2E). Like SOX9 progenitors, *Cebpa* mutant cells were also much more proliferative (KI67+ in 25% 3414 GFP+ cells from 3 mice, compared to 6% 2737 GFP+ cells from 3 mice in the control), likely contributing to their clustering (Fig. 2E, 2F). Therefore, without CEBPA, neonatal AT2 cells reduce their AT2 program and gain plasticity toward SOX9 progenitors and, to a much lesser extent, AT1 cells.

### Single-cell multiome defines CEBPA-dependent neonatal AT2 cell program and plasticity

To fully characterize the CEBPA-dependent changes, we FASC purified E-Cadherin+ epithelial cells from our neonatal *Cebpa* mutant and littermate control lungs and performed single-cell multiome for concurrent profiling of their transcriptomes and epigenomes (Fig. S3A). On the combined single-cell UMAPs, club, ciliated, and AT1 cells from control and mutant lungs formed superimposed clusters, suggesting minimal changes, as they were not targeted by *Sftpc^CreER^* and lacked the GFP transcript from *Rosa^Sun1GFP^* (Fig. 3A, 3B, S3B). In contrast, while 3.7% (267 out of 7047 cells) AT2 cells from the mutant lung were intermixed with those from the control and still expressed *Cebpa*, consistent with them being escapers of deletion, the rest formed a separate *Cebpa*^−^ cluster (mutant AT2), as well as a proliferative cluster mainly made of cells from the mutant lung as predicted by KI67 immunostaining (Fig. 3A, 3B). A marker *Lamp3* and a 100-gene signature score for AT2 cells were reduced in mutant AT2 cells, whereas a marker *Sox9* and a 119-gene signature score for SOX9 progenitors were increased (Fig. 3C). The mutant AT2 cluster extended toward the AT1 cell cluster, forming a bridge that was GFP+ and thus descendant of recombined AT2 cells (Fig. 3C). This bridging population was specific to the mutant lung and expressed a marker *Hopx* and a 100-gene signature for AT1 cells, but still clustered separately from normal AT1 cells, possibly due to their AT2 cell origin and limited time for AT1 differentiation after *Cebpa* deletion. Considering the observed HOPX immunostaining in LAMP3-non-cuboidal cells (Fig. S3C, 2D), we named this bridging population HOPX+ AT1-like cells. Differential expression analysis of *Cebpa* mutant versus control AT2 cells confirmed downregulation of AT2 genes (e.g. *Lyz2*, *Lyz1*, *Sftpb*, and *Il33*) and upregulation of progenitor (e.g. *Sox9*, *Clu*, and *Col18a1*) and AT1 genes (e.g. *Akap5*, *Fbln5*, and *Rtkn2*), although some surfactant genes including *Sftpc* were less reduced, possibly due to RNA perdurance or redundant transcriptional activation (Fig. 3D, S3D, S3E).

**Figure 3.**
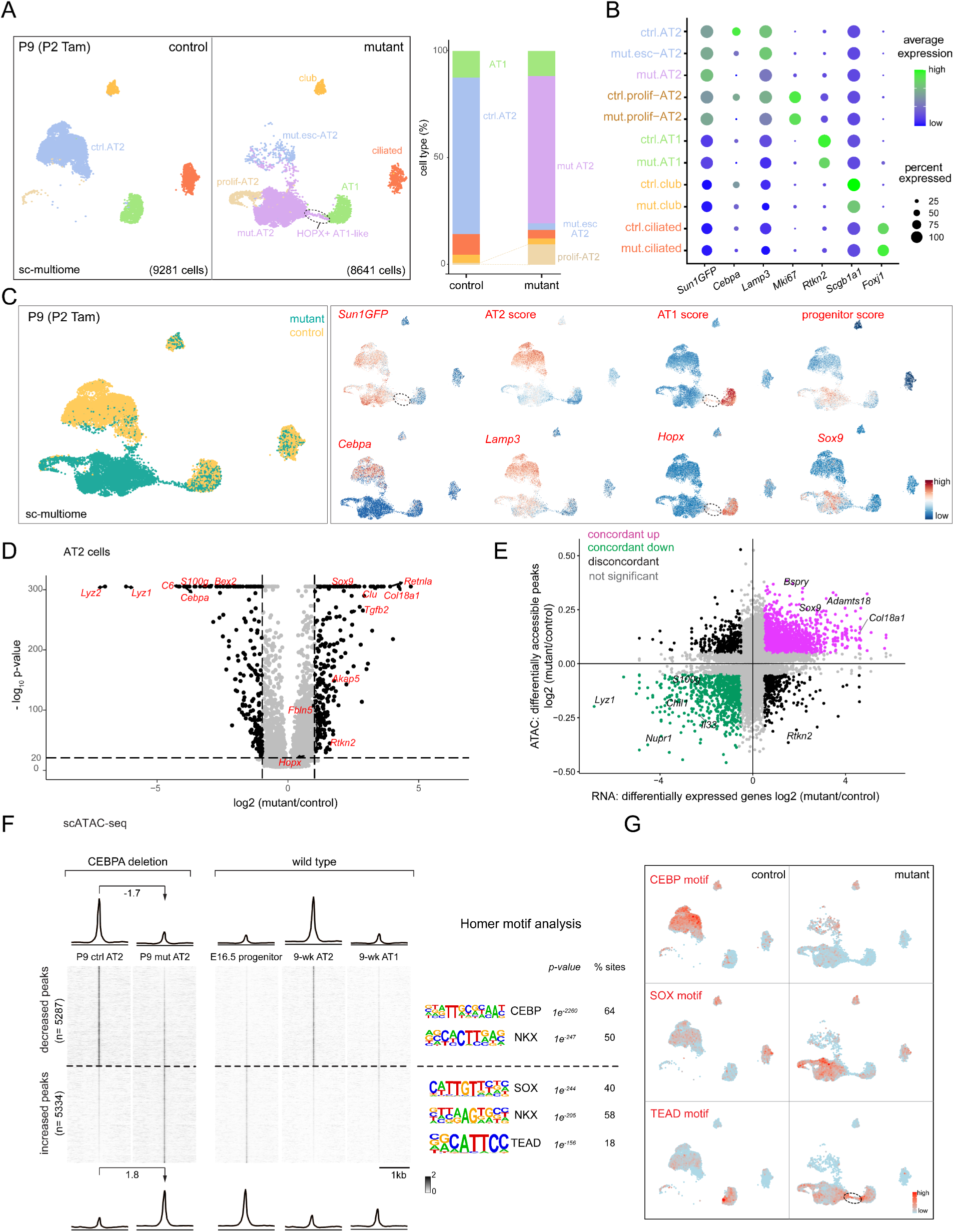
Single-cell multiome defines CEBPA-dependent neonatal AT2 cell program and plasticity. (**A**) Sc-multiome UMAPs of purified epithelial cells from *Cebpa* mutant and littermate control lungs color coded by cell type (left) and the corresponding percentages (right). Esc, escaper; prolif, proliferating; Tam, 250 ug tamoxifen. See Fig. S3A for sorting strategy. (**B**) Dot plot showing the lineage marker (Sun1GFP), *Cebpa*, and cell type markers. See also feature plots in (**C**) and Fig. S3B. (**C**) Sc-multiome UMAP color coded for genotype (left) and feature plots of metagene scores (top) and representative genes (bottom). Circled population is specific to the mutant, GFP+, and expresses AT1 genes, thus labeled as HOPX+ AT1-like cells in (**A**). See Table S3 for metagene lists. (**D**) Volcano plot showing downregulation of AT2 genes and upregulation of progenitor genes in mutant AT2 cells compared to control AT2 cells defined in (**A**). (**E**) Scatter plot correlating changes in the accessibility of scATAC-seq peaks (y-axis) and scRNA-seq expression of their nearest genes (x-axis), color coded as concordant or discordant as well as the directionality of change. (**F**) ScATAC-seq heatmaps and profile plots of decreased and increased peak sets in the mutant and associated log2 fold changes, as well as the corresponding scATAC-seq data in wild type cells and associated Homer motifs. (**G**) Feature plots of motif activity scores showing that the mutant has lower CEBP, higher SOX and TEAD (circle) activities. See Table S3 for raw data.

To explore the epigenetic mechanism of the transcriptional changes, we performed differential accessibility analysis of *Cebpa* mutant versus control AT2 cells as pseudobulks and identified 10,621 differential peaks. Assigning each peak to its nearest gene, we found a high concordance (71%) between peaks and gene expression, with increases in both (39%) for progenitor genes including *Sox9*, *Bspry*, and *Adamts18* and decreases in both (32%) for AT2 genes including *Lyz1*, *Il33*, and *S100g* (Fig. 3E). Moreover, the 5,287 decreased peaks were normally more accessible in AT2 cells, in comparison to AT1 and SOX9 progenitor cells, and were enriched for CEBP and NKX motifs, consistent with *Cebpa* deletion (Fig. 3F, 3G). The 5,334 increased peaks were normally more accessible in SOX9 progenitor or, to a lesser extent as expected from the small number of HOPX+ AT1-like cells, AT1 cells, in comparison to AT2 cells, and were enriched for SOX and TEAD motifs, consistent with activation of progenitor and AT1 programs (Fig. 3F, 3G). Therefore, besides the more predictable role of CEBPA in promoting the AT2 program, marker and whole-genome analyses unexpectedly show that neonatal AT2 cells have the plasticity to revert to SOX9 progenitors when unconstrained by CEBPA. The 5,334 increased peaks represent CEBPA-dependent plasticity of neonatal AT2 cells that will be explored later.

### CEBPA recruits NKX2-1 to promote the AT2 program and indirectly restricts the progenitor program

The identification of differential accessibility peaks that were largely concordant with differential gene expression prompted mechanistic analysis to link ATAC peaks to CEBPA chromatin binding. Given the enrichment in CEBP and NKX motifs (Fig. 3F) and the normal expression of NKX2-1 in the *Cebpa* mutant (Fig. S3F), we used our published cell-type-specific ChIP-seq protocol^8^ to perform CEBPA ChIP-seq on control AT2 cells at P2, the time of Cre-recombination, as well as NKX2-1 ChIP-seq on AT2 cells from control and *Cebpa* mutant lungs at P8 (Fig. S4A, 4A). The 5,287 decreased peaks, as exemplified by a peak near an AT2 gene *Il33*, were bound by CEBPA and NKX2-1; NKX2-1 binding decreased upon *Cebpa* deletion, suggesting recruitment of NKX2-1 by CEBPA. NKX2-1 binding at these sites was specific to AT2 cells but not E14.5 SOX9 progenitors, suggesting that CEBPA was required to acquire AT2-specific NKX2-1 binding (Fig. 4A, 4B). The predicted CEBP and NKX motifs, but not SOX motif, were concentrated at the center of the decreased peaks (Fig. 4A). Furthermore, the average distance between CEBPA and NKX2-1 binding sites were 54 bp, consistent with proximity or even direct binding between CEBPA and NKX2-1, although such protein-protein interaction needed technically challenging biochemical studies of purified AT2 cells.

**Figure 4.**
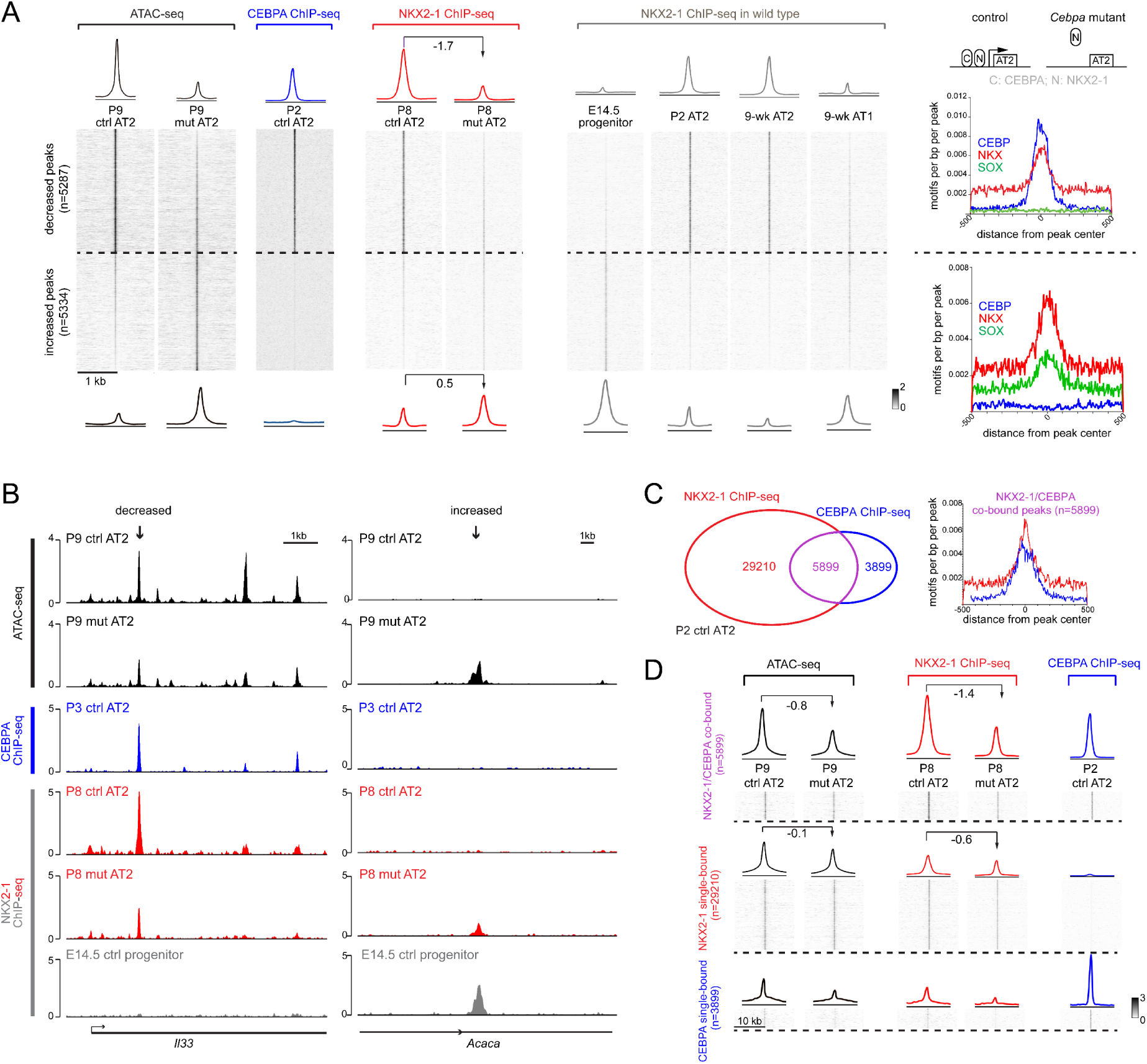
CEBPA recruits NKX2-1 to promote the AT2 program and indirectly restricts the progenitor program. (**A**) Heatmaps and profile plots of CEBPA and NKX2-1 binding for decreased and increased peak sets from Fig. 3F, as well as associated frequency distributions of CEBP, NKX, SOX motifs. CEBPA binds to decreased peaks but not increased peaks. NKX2-1 binding decreases (log2 fold change) for decreased peaks and increases (log2 fold change) for increased peaks in the mutant, corresponding to AT2 and progenitor/AT1-specific binding in wild type lungs. Inset: a recruitment model, in which CEBPA normally recruits NKX2-1 to activate AT2 genes, whereas without CEBPA, NKX2-1 is released from AT2 genes and possibly relocates to progenitor and AT1 genes. See Fig. S4A for nuclei sorting strategy. (**B**) Representative coverage plots of (**A**) showing a decreased peak near an AT2 gene *Il33*, and an increased peak near a progenitor gene *Acaca*. (**C**) Venn diagram showing NKX2-1 and CEBPA co-bound and single-bound peak sets in purified P2 AT2 cells (left) and frequency distributions of NKX and CEBP motifs for the co-bound peak set (right). (**D**) Heatmaps and profile plots for the 3 peak sets in (**C**) and associated log2 fold changes showing largest decreases for the co-bound peak set. See Table S4 for raw data.

In contrast, the 5,334 increased peaks, as exemplified by a peak near a progenitor gene *Acaca*, had little CEBPA binding and a slight increase in NKX2-1 binding upon *Cebpa* deletion, possibly attributable to its relocation to progenitor and AT1-specific sites without sequestration by CEBPA (Fig. 4A, 4B). Interestingly, NKX2-1 bound more at these sites in E14.5 SOX9 progenitors as well as AT1 cells compared to AT2 cells, suggesting that these progenitor-specific NKX2-1 binding sites normally lost NKX2-1 binding and were closed during alveolar differentiation, but were reopened upon *Cebpa* deletion (Fig. 4A). Alternatively, given the robust SOX9 expression and the prevalent SOX motif for these sites (Fig. 2, 3), CEBPA directly or indirectly repressed *Sox9*, which in turn initiated the progenitor program. Indeed, we identified a putative regulatory region 3’ to *Sox9* that was open in the SOX9 progenitors and reopened in the *Cebpa* mutant, a profile mirrored by NKX2-1 binding (Fig. S4B).

The link between CEBPA/NKX2-1 chromatin binding and differential accessibility peaks was also examined in the reverse direction. CEBPA and NKX2-1 binding sites in AT2 cells were categorized as co-bound and single-bound for each TF (Fig. 4C). Compared to NKX2-1 single-bound sites, CEBPA/NKX2-1 co-bound sites, had a greater decrease in NKX2-1 binding and accessibility upon *Cebpa* deletion, reinforcing the said recruitment model and implicating other regulators of NKX2-1 binding and accessibility at the NKX2-1 single-bound sites (Fig. 4D). The CEBPA single-bound sites had limited accessibility and NKX2-1 binding as well as limited changes, suggesting a minor impact of CEBPA on its own (Fig. 4D). Taken together, in AT2 cells, CEBPA recruits NKX2-1 to promote the AT2 program whereas CEBPA-dependent plasticity sites are not bound by CEBPA.

### CEBPA maintains the AT2 program without affecting the progenitor program in mature AT2 cells

As the transcriptomic and epigenomic landscape of AT2 cells matured postnatally (Fig. 1), we posited that they would reinforce their gene regulatory network and exhibit less cell plasticity. To test this, we induced Cre-recombination in mature AT2 cells in >5-week old lungs and achieved 92% efficiency in deleting *Cebpa* (1919 GFP+ cells from 3 mice), again without affecting CEBPA expression in alveolar macrophages (Fig. 5A). As in the neonatal deletion model, mature *Cebpa* mutant AT2 cells downregulated LAMP3, lost IL33, and had fewer lamellar bodies (Fig. 5A, 5B, S4C, S4D). However, they did not express SOX9 or KI67, suggesting a loss of cell plasticity toward SOX9 progenitors (Fig. 5A).

**Figure 5.**
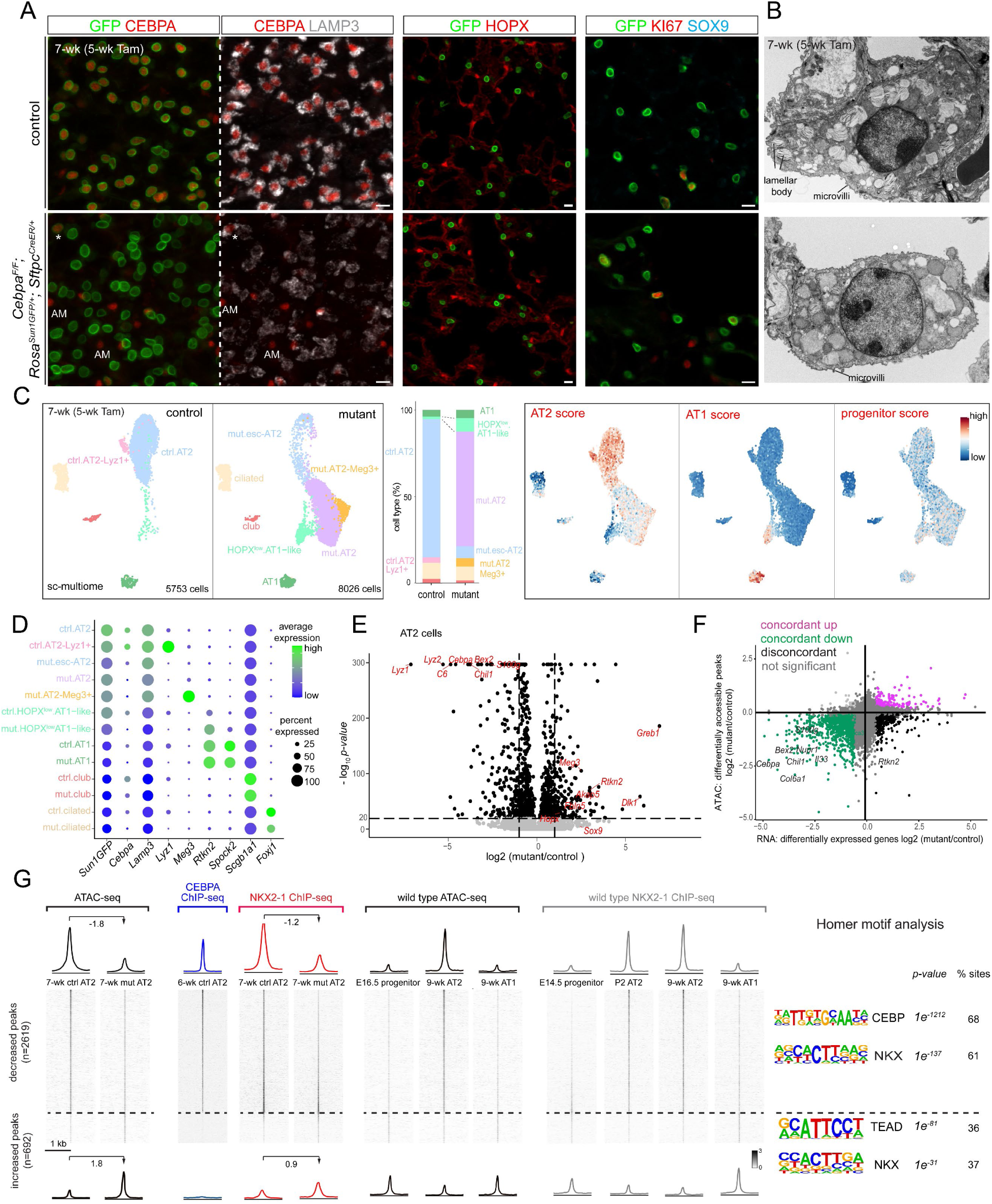
CEBPA maintains the AT2 program without affecting the progenitor program in mature AT2 cells. (**A**) Confocal images of immunostained adult AT2-specific *Cebpa* mutant and littermate control lungs showing loss of CEBPA in GFP+ recombined cells (asterisk: escaper), without affecting its expression in alveolar macrophages in the airspace (AM), and reduced LAMP3, but no extra HOPX, SOX9, or KI67. Tam, two doses of 3 mg each tamoxifen at 48 hr interval (same for the rest of Fig. 5). Scale: 10 um. (**B**) TEM images showing reduction in lamellar bodies in mutant AT2 cells without affecting their apical microvilli. Large granules in mutant AT2 cells lack characteristic lamellae. Scale: XX um. See Fig. S4D for quantification. (**C**) Sc-multiome UMAPs of purified epithelial cells from *Cebpa* mutant and littermate control lungs color coded by cell type (left), the corresponding percentages (middle), and metagene scores. Esc, escaper. See Table S3 for metagene lists. (**D**) Dot plot showing the lineage marker (Sun1GFP), *Cebpa*, and cell type markers. *Rtkn2*, but not *Spock2*, is expressed in HOPX_low_ AT1-like cells. (**E**) Volcano plot showing downregulation of AT2 genes but minimal upregulation of progenitor/AT1 genes in mutant AT2 cells compared to control AT2 cells defined in (**C**). Compare with Fig. 3D. (**F**) Scatter plot correlating changes in the accessibility of scATAC-seq peaks (y-axis) and scRNA-seq expression of their nearest genes (x-axis), color coded as concordant or discordant as well as the directionality of change. Compared to Fig. 3E, few concordant pairs are upregulated. See Table XX for the complete list. (**G**) Heatmaps and profile plots of decreased and increased scATAC-seq peak sets in the adult mutant and associated log2 fold changes, as well as the corresponding CEBPA and NKX2-1 binding and scATAC-seq data in wild type cells and associated Homer motifs. Decreased peaks have CEBPA binding and decreased NKX2-1 binding, corresponding to ATAC accessibility and NKX2-1 binding in wild type AT2 cells. Increased peaks are many fewer and have no CEBPA binding and increased NKX2-1 binding, corresponding to NKX2-1 binding in wild type AT1 cells. See Table S5 for raw data.

Single-cell multiome profiling of E-Cadherin+ epithelial cells from mature *Cebpa* mutant and littermate control lungs showed a transcriptional shift only in targeted GFP+ AT2 cells (Fig. 5C, 5D). Notwithstanding additional heterogeneity including a *Lyz1*+ population in the control lung and a *Meg3*+ population in the mutant, possibly related to lung cancer and fibrosis^12,13^, the most prominent change was downregulation of AT2 genes in *Cebpa*^−^ mutant AT2 cells, without activating progenitor genes or forming a proliferative population as in the neonatal lungs (Fig. 5C). Compared to the control lung, the *Cebpa* mutant lung had a larger population clustered near AT2 cells and expressing some but not all AT1 gene transcripts (Fig. 5C, 5D), although few HOPX+ cells were detected by immunostaining (Fig. 5A). Accordingly, we considered this population HOPX^low^ AT1-like cells to indicate their limited AT1 differentiation (Fig. 5C). The reduction in the AT2 program, no increase in the progenitor program and limited increase in the AT1 program were supported by differential gene expression analysis (Fig. 5E).

Compared to the neonatal model, differential accessibility analysis of *Cebpa* mutant versus control mature AT2 cells identified 2,619 decreased peaks but only 692 increased peaks, suggesting less CEBPA-dependent cell plasticity than neonatal AT2 cells. These differential peaks were still 76% concordant with gene expression (Fig. 5F). The decreased peaks were AT2-specific, enriched for CEBP and NKX motifs, had CEBPA and NKX2-1 binding in control AT2 cells but decreased NKX2-1 binding in *Cebpa* mutant AT2 cells, and had NKX2-1 binding in normal AT2 but not progenitor nor AT1 cells, supporting the same recruitment model of NKX2-1 by CEBPA in mature AT2 cells (Fig. 5G, S4E, 4A inset). The few increased peaks had no CEBPA binding, were enriched for NKX and TEAD motifs, but not SOX motif, and had some accessibility enriched for progenitors and AT1 cells but to a much lesser extent than the neonatal increased peaks (Fig. 5G, S4E). NKX2-1 binding in purified control AT2 cells was low but increased in *Cebpa* mutant AT2 cells, possibly due to its limited redistribution to AT1-specific sites in the considerable number of HOPX^low^ AT1-like cells (Fig. 5G).

The main difference between the mature versus neonatal *Cebpa* deletion models was the inability of mature AT2 cells to reactive the SOX9 progenitor program. This decrease in cell plasticity as AT2 cells matured were molecularly defined as the CEBPA-dependent, increased peaks unique to neonatal AT2 cells (5,124 peaks in Fig. S4F). The neonatal-specific plasticity was in regions that were accessible in progenitors and closed for 2 days versus 35 days when Cre-recombination was induced in neonatal versus mature lungs, respectively (Fig. S4F). The duration of chromatin closure might lead to less reversible changes in histone modifications, DNA methylation, or high-order chromatin structure across the sites of differential plasticity or a few nodal sites of master genes, such as *Sox9*. Although CEBPA did not bind to these differentially plastic sites, its deletion revealed their presence.

Taken together, in neonatal and mature AT2 cells, CEBPA recruits NKX2-1 to promote and maintain the AT2 program; without CEBPA, neonatal but not mature AT2 cells have the plasticity to reactivate the SOX9 progenitor program.

### Viral infection expands CEBPA-dependent plasticity in mature AT2 cells

The temporal restriction in cell plasticity from neonatal to mature AT2 cells reminded us of the doctrine that injury-repair recapitulates development and prompted us to test if respiratory virus infection would reactivate the neonatal plasticity in mature AT2 cells. We infected our mature *Cebpa* deletion model with Sendai virus, which was known to preferentially injure AT2 cells, forming AT2-less regions, and trigger AT2 cell proliferation 14 days post infection^14^. Strikingly, while the infected control lung repaired itself with no SOX9 expression and only isolated KI67 expression in AT2 cells, the infected *Cebpa* mutant lung had SOX9 expression in 8% of AT2 cells (7776 GFP+ cells from 3 mice) and clusters of KI67+ AT2 cells, reminiscent of the neonatal *Cebpa* mutant (Fig. 6A). SOX9+ AT2 cells often abutted lobe edges or airways and macro-vessels, topologically distal ends of the respiratory tree favoring de novo growth as we described^14^ (Fig. S5A). The regional preference, in conjunction with localized virus delivery, suggested that the percentage of mutant AT2 cells capable of expressing SOX9 could be much higher. SOX9 activation depended on infection because saline treated control and *Cebpa* mutant lungs did not express SOX9 (Fig. S5B).

**Figure 6.**
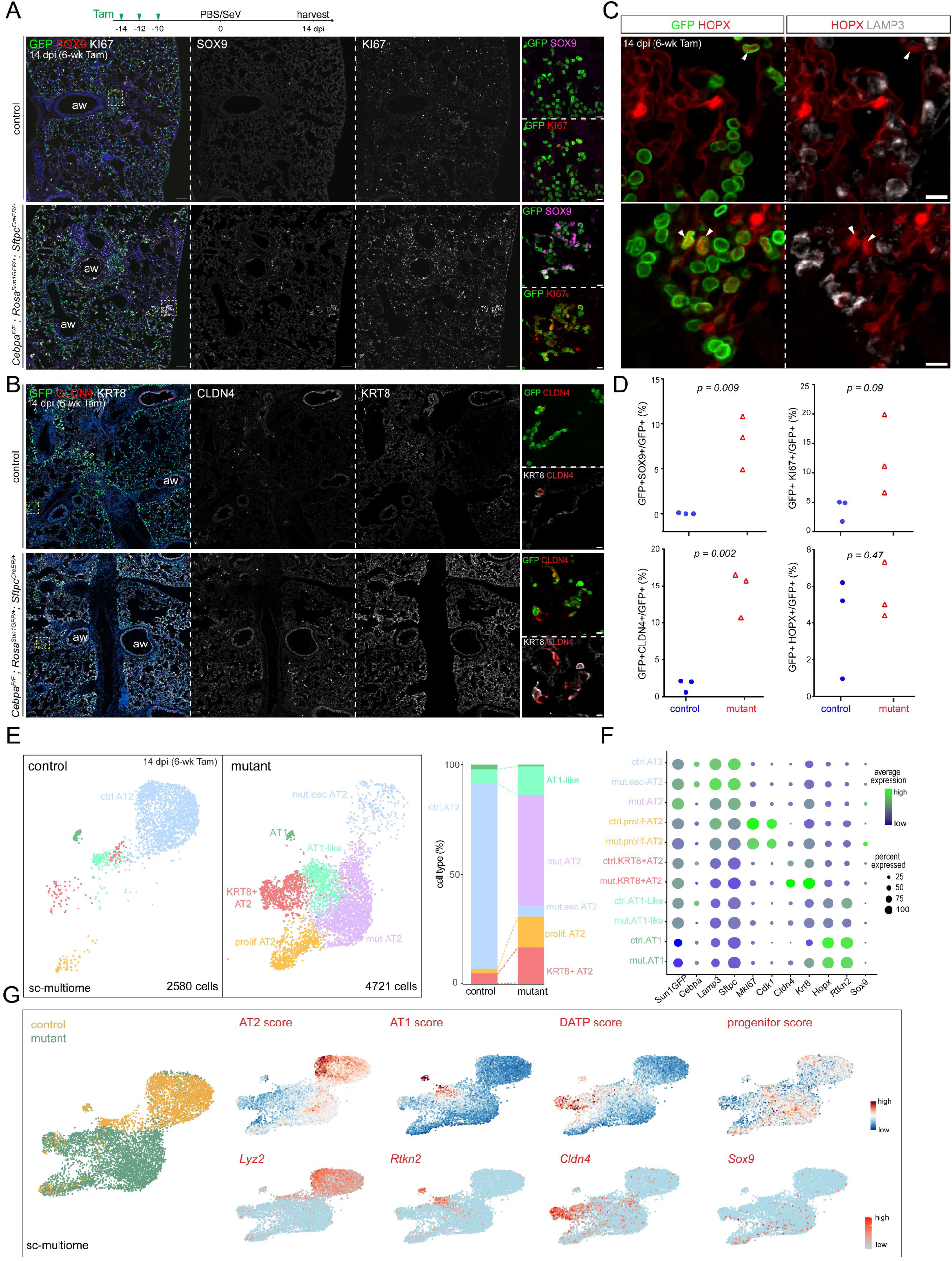
Viral infection expands CEBPA-dependent plasticity in mature AT2 cells. (**A**) Experimental timeline of tamoxifen injection (Tam, 3 mg), Sendai virus (SeV) or saline (PBS) administration, and lung harvest at 14 dpi (day post-infection). Confocal images of immunostained infected *Cebpa* mutant and littermate control lungs, showing mutant-specific activation of SOX9 and increase in KI67 near airways (aw) and lobe edges (inset; scale: 10 um). Scale: 100 um. (**B**) Confocal images of lungs in (**A**) showing increased KRT8 and CLDN4. Scale: 100 um (inset: 10 um). (**C**) Confocal images of lungs in (**A**) showing lineage-labeled HOPX+ cells with little LAMP3 (arrowhead). Scale: 10 um. (**D**) Quantification of (**A**), (**B**) and (**C**). Each symbol represents one mouse from littermate pairs (Student’s t-test). (**E**) Sc-multiome UMAPs of purified epithelial cells from infected *Cebpa* mutant and littermate control lungs color coded by cell type (left) and the corresponding percentages (right). Esc, escaper; prolif, proliferating. (**F**) Dot plot showing the lineage marker (Sun1GFP), *Cebpa*, and cell type markers. (**G**) Sc-multiome UMAP color coded for genotype (left) and feature plots of metagene scores (top) and representative genes (bottom). A published damage-associated transient progenitor (DATP) score_18_ marks KRT8/CLDN4+ cells. See Table S6 for raw data including metagene lists.

E-Cadherin+ epithelial cells from infected control and *Cebpa* mutant lungs were profiled with single-cell multiome (Fig. 6E, 6F). As in prior neonatal and mature mutant models, *Cebpa*-AT2 cells from the infected mutant lung clustered separately from escapers of deletion, as well as AT2 cells in the infected control lung. *Sox9* and progenitor signature were higher in the mutant, albeit to a lesser extent due to the said spatial restriction (Fig. 6G). Proliferative AT2 cell cluster was much more prominent in the mutant, corroborating the KI67 immunostaining, and expressed *Sox9*, suggesting a possible coupling between cell cycle and SOX9 activation (Fig. 6F, 6G).

Sendai virus infection also induced in both control and mutant lungs two other GFP+ AT2-derived cell populations: KRT8/CLDN4+ transitional cells and AT1-like cells, marked by respective gene signatures (Fig. 6G). By immunostaining, the former had high KRT8 and ectopic CLDN4, but low LAMP3 and no HOPX; the latter had HOPX but no LAMP3 and were no longer cuboidal (Fig. S5C, S5D). Although the two populations could represent sequential steps during AT2 to AT1 differentiation^15–18^, their locations on the UMAPs were also compatible with two parallel states with only the AT1-like cells transitioning to AT1 cells and KRT8/CLDN4+ cells being arrested. Regardless, the *Cebpa* mutant lungs had a dramatic expansion of KRT8/CLDN4+ transitional cells, as confirmed by immunostaining, which were distinct from SOX9-expressing cells (Fig. 6B, S6A). Despite the higher number of AT1-like cells captured by single-cell multiome, they were not reliably detected by HOPX immunostaining, possibly due to the higher sensitivity of single-cell multiome in documenting the gradual AT2-AT1 transition (Fig. 6C, 6D, 6E). CEBPA was normally lost in both populations even in the control lung by single-cell profiling and immunostaining (Fig. S6B, S6C), consistent with their decreased/lost LAMP3 expression and the described role of CEBPA in maintaining the AT2 program. Therefore, the population expansion in the mutant was likely because most AT2 cells became eligible for alternative fates as the result of losing CEBPA. Furthermore, as the loss of CEBPA in KRT8/CLDN4+ and AT1-like cells alone was insufficient to activate *Sox9* in the infected control lung, SOX9 activation in the infected mutant represented a separate plasticity from injury-induced loss of the AT2 program and adoption of KRT8/CLDN4+ and AT1-like programs. Taken together, Sendai virus infection increases the plasticity of mature AT2 cells, which manifests upon *Cebpa* deletion as activation of the SOX9 progenitor program and expansion of the KRT8/CLDN4+ program.

## DISCUSSION

In this study, we have tracked the transcriptomic and epigenomic changes as well as plasticity of AT2 cells during their specification and subsequent maturation and upon viral injury. We show that the AT2-specific TF CEBPA recruits the lung lineage TF NKX2-1 to promote and maintain the AT2 program. *Cebpa* deletion also reveals an evolving landscape of AT2 cell plasticity shaped by the developmental history and external stimuli. In wild type lungs, neonatal and mature AT2 cells form AT1-like cells infrequently during homeostasis, but more readily upon viral injury and additionally form KRT8/CLDN4+ transitional cells (Fig. 7A). Without CEBPA, these processes are enhanced; most strikingly, neonatal but not mature AT2 cells activate the SOX9 progenitor program and this neonatal plasticity is bestowed to mature AT2 cells by viral injury (Fig. 7A). The plasticity landscape is submerged by the AT2 program, partially exposed when altered sufficiently, but unmasked upon *Cebpa* deletion (Fig. 7B). This interplay between the sea-level program and submerged plasticity landscape could be important in generating and engrafting the most effective lung cells in cell therapy and in activating endogenous stem cells without forming a tumorigenic landscape.

**Figure 7.**
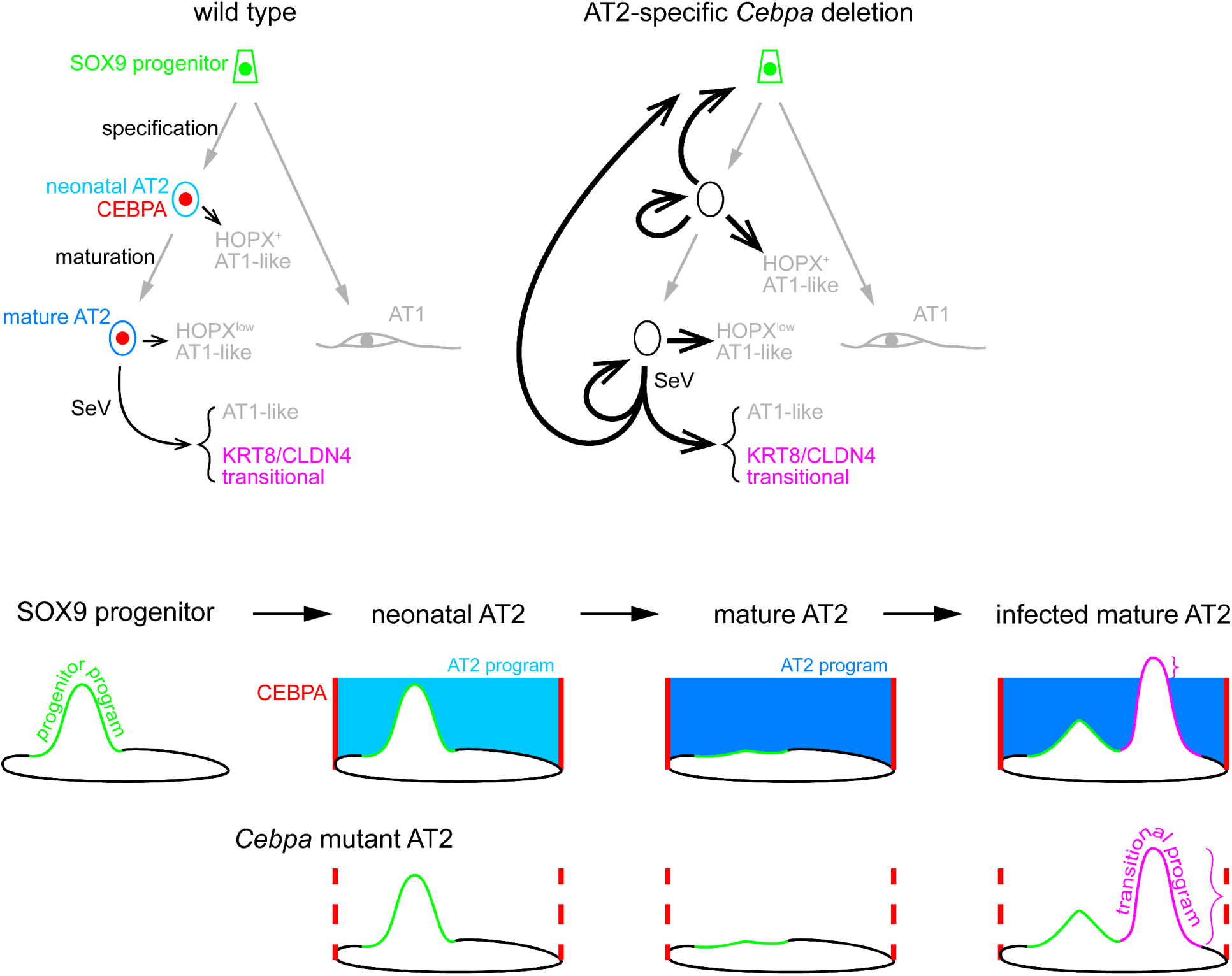
Diagram of CEBPA restricting AT2 cell plasticity during development and injury-repair. Top: in a wild type lung, SOX9 progenitors undergo specification and maturation to become neonatal/nascent and mature AT2 cells, sequentially, expressing CEBPA. A small fraction of AT2 cells at both stages become AT1-like cells with varying levels of HOPX. Sendai virus (SeV) infection induces KRT8/CLDN4 transitional cells as well as AT1-like cells, as a sequential or parallel response of injury-repair. *Cebpa* deletion in neonatal, but not mature, AT2 cells leads to reversion to the progenitor program and proliferation, in addition to an increase of AT1-like cells upon either deletion. Sendai virus infection bestows mature AT2 cells with the plasticity to revert to progenitors as well as to more readily transition to the KRT8/CLDN4 state. Bottom: in an aerial view of the cell plasticity landscape, the progenitor program, although still present in neonatal AT2 cells, is submerged under the AT2 program promoted by CEBPA. Without CEBPA, the AT2 program subcedes to expose the progenitor program. As AT2 cells mature, the progenitor program disappears so that, even without the AT2 program, no progenitor program is visible. After infection, mature AT2 cells reshape their plasticity landscape, but only the KRT8/CLDN4 transitional program rises enough to manifest over the AT2 program. Without CEBPA and the AT2 program, the progenitor program is visible and the transitional program more evident.

A gene regulatory network of alveolar cell fates is emerging from recent mechanistic studies of alveolar TFs^8,19^. Both AT1 and AT2 cells express the lung lineage TF NKX2-1 and require NKX2-1 to express AT1 or AT2-specific genes, respectively, and to suppress gastrointestinal (GI) genes. The cell-type-specific functions of NKX2-1 are attributed to its cell-type-specific chromatin binding, as the result of its recruitment by cell-type-specific co-TFs: YAP/TAZ/TEAD in AT1 cells and CEBPA in AT2 cells^8^. This symmetry between AT1 and AT2 cells is skewed with respect to cell plasticity. Neonatal *Yap/Taz* mutant AT1 cells readily become AT2-like, whereas only 12% neonatal *Cebpa* mutant AT2 cells become AT1-like with the vast majority (80%) activating the SOX9 progenitor program (Fig. 2). Perhaps AT2 cells are more closely related to progenitors in gene expression and cell morphology than AT1 cells are, as possibly needed for their facultative stem cell function. Alternatively but compatibly, AT2 differentiation might be the default path of SOX9 progenitors whereas AT1 differentiation requires YAP/TAZ activation by mechanical stretching in the growing neonatal lung. As a result, *Yap/Taz* mutant AT1 cells fail to relay the mechanical signal to AT1-specific NKX2-1 recruitment and default to AT2 differentiation, whereas *Cebpa* mutant AT2 cells, surrounded by normal AT1 cells, are not subject to mechanical stretching and AT1 differentiation. The neonatal plasticity becomes limited in the mature lung where *Yap/Taz* mutant AT1 cells less readily become AT2-like and *Cebpa* mutant AT2 cells do not become SOX9 progenitors^20^. Sendai virus injury reactivates the SOX9 progenitor program in *Cebpa* mutant AT2 cells, whereas hyperoxia injury or blocking mechanical stretching stimulates the AT2 program even in wild type AT1 cells, although more studies are needed to understand the predicted shedding of excessive cell membrane during AT1-to-AT2 conversion^20,21^. Future mechanistic studies are expected to expand the core NKX2-1/YAP/TAZ/TEAD/CEBPA network to include additional TFs, such as KLF5 and AP-1 members, and effector enzymes, such as histone modifiers and chromatin remodelers^22^.

This study illustrates an experimental and conceptual framework of cell plasticity and highlights three associated challenges: its diversity, definition, and regulation. First, cell plasticity can be forced through overexpression of TFs during cell reprogramming in culture or in vivo^1,23^. Although cell plasticity from gene deletion is likely more physiological, both deleting and overexpressing different TFs in the same cell can lead to different plasticity, as seen in the AT1-AT2-progenitor plasticity triad from targeting YAP/TAZ/CEBPA versus the lung-GI dyad from targeting NKX2-1 (this study and ^8,19^). Instead of submitting to the notion of universal plasticity from arbitrary use of the same DNA blueprint, we need systematic precise gene perturbation coupled with quantitative molecular readouts of cell plasticity.

Relatedly, the second challenge arises from qualitative, categorical, or at times semantic definitions of a cell type. Genome-wide profilings have revealed molecular variants of classical cell types, invoking prefixes such as pro-, pre-, semi-, and quasi- and postfixes such as -like and -oid, as well as adjectives such as intermediate, transitional, and hybrid – several of which are also used in this study. As experimental and computational technology advances, cell types and states will be objectively digitized and their changes upon perturbations become the definition of cell plasticity. Our use of the accessibility changes upon *Cebpa* deletion in neonatal or mature AT2 cells is such an attempt. A better definition of cell type and cell plasticity will also help clarify comparable terms in the literature including cell potency, lineage infidelity, and points of no return^24–26^.

Last, although we show that CEBPA restricts and its deletion reveals AT2 cell plasticity, the molecular correlates and regulators of the shifting plasticity landscape are unclear. Sites of plasticity may be primed/poised via bivalent histone marking, opened de novo by pioneering TFs, or reset upon DNA/histone synthesis during cell cycle^27–29^. Regardless, CEBPA does not bind to plasticity sites, as in the lack of binding of NKX2-1 to GI genes^19^. The recruitment model predicts free diffusion and ectopic binding of the partner TF in the absence of the recruiting TF, such that FOXA2 might activate GI genes without NKX2-1 and NKX2-1 might activate AT2 genes without YAP/TAZ^8,30^. However, there is limited gain of NKX2-1 binding at the plasticity sites in both neonatal and mature *Cebpa* mutant AT2 cells (Fig. 4, 5). Rather than redistributing CEBPA’s partner TF NKX2-1, the neonatal *Cebpa* mutant cells transcriptionally activate SOX9, whose motif causally or coincidentally dominates the plasticity sites. These plasticity sites, such as the one near *Sox9*, are open in SOX9 progenitors several days prior and become closed and unresponsive to *Cebpa* deletion in mature AT2 cells, suggesting a role of developmental history in shaping cell plasticity. Reactivation of developmental plasticity upon viral injury (Fig. 6), or possibly during tumorigenesis, may rewrite cells’ developmental history and represent therapeutic opportunities.

### Limitations of study

This study has not explored the translational significance of AT2 cell plasticity nor extended our findings to the human lung. It is unclear if CEBPA-dependent AT2 cell plasticity is reactivated in response to chemical and biological injuries other than Sendai virus. In addition, our focused mutant models of CEBPA does not test the causal roles of other TFs including SOX9 and NKX2-1 in the *Cebpa* mutant, nor the transcriptional control of AT2-specific expression of *Cebpa*.

## METHODS

### Mice

*Cebpa^F^* ^31^ were obtained from the Jackson Laboratory (stock #006447). *Sftpc^CreER^* ^32^ and *Rosa^Sun1GFP^* ^33^ have been previously described. Mice were given intraperitoneal injection of tamoxifen (T5649, Sigma) dissolved in corn oil (C8267, Sigma). The doses and time of injection were described in figure legends. All mice were housed in the MD Anderson facility and the proposed studies will be performed in accordance with all federal regulations on the use of animals in research and have been approved by the Institutional Animal Care and Use Committee at the University of Texas MD Anderson Cancer Center.

### Antibodies

For immunofluorescence, the following antibodies were used: rabbit anti-CCAAT/enhancer binding protein alpha (C/EBPA, 1:500, 8178P, Cell Signaling Technology), chicken anti-green fluorescent protein (GFP, 1:5000, AB13970, Abcam), rabbit anti-NK homeobox 2-1 (NKX2-1, 1:1000, sc-13040, Santa Cruz), rabbit anti-pro-surfactant protein C (SFTPC, 1:1000, AB3786, Millipore), goat anti-SOX9 (SOX9, 1:1000, AF3075, R&D Systems), rabbit ani-SOX9 (SOX9, 1:1000, AB5535, Millipore). Goat Anti-Mouse Il-33 (IL33, 1:500, R&D, AF3626). rabbit anti-homeodomain only protein (HOPX, 1:500, sc-30216, Santa Cruz). mouse anti-homeodomain only protein (HOPX, 1:250, sc-398703 AF647, Santa Cruz). rat anti-KI67 (KI67, 1:1000, 14-5698-82, Invitrogen), guinea pig anti-lysosomal associated membrane protein 3 (LAMP3, 1:500, 391005, SySy), rat anti-epithelial cadherin (ECAD, 1:1000, 13190, Invitrogen).

The following antibodies were used for FACS: PE/Cy7 rat anti-CD45 (CD45, 1:250, 103114, BioLegend), PE rat anti-epithelial cadherin (ECAD, 1:250, 147304, BioLegend), BV421 rat anti-epithelial cell adhesion molecule (EPCAM, 1:250, 118225, BioLegend), and AF647 rat anti-Intercellular adhesion molecule 2 (ICAM2, 1:250, A15452, Thermo Fisher).

The following antibodies were used for chromatin immunoprecipitation: rabbit anti-NK Homeobox 2-1 (NKX2-1, 1 µg per reaction, ab133737, Abcam) and rabbit anti-CEBPA (C/EBPα, (D56F10) XP, 1 µg per reaction, Cell Signaling Technology).

### Section immunofluorescence

After intraperitoneal injection of Avertin (Sigma, T48402), the heart’s right ventricle was subjected to perfusion using phosphate-buffered saline (PBS, pH 7.4). Subsequently, the trachea was cannulated and the lungs were inflated with a solution of 0.5% paraformaldehyde (Sigma, P6148) in PBS, maintaining a pressure of 25 cm H_2_O. The lungs were fixed in 0.5% PFA in PBS for 3-4 hr at room temperature and then washed with PBS overnight at 4°C. For section immunostaining, the fixed lung lobes were cryoprotected overnight at 4°C in 20% sucrose in PBS containing 10% optimal cutting temperature compound (OCT; 4583, Tissue-Tek, Tokyo, Japan; optional for embryonic lungs) and then embedded in OCT and frozen in −80°C. Immunostaining of frozen sections was adapted from previously described method^34^. Briefly, frozen sections at 10-μm thickness were blocked in PBS with 0.3% Triton X-100 and 5% normal donkey serum (017-000-121, Jackson ImmunoResearch) and then incubated with primary antibodies diluted in PBS with 0.3% Triton X-100 at 4°C overnight in a humidified chamber. The following day, the sections were washed with PBS for 1 h at room temperature and then incubated with secondary antibodies (Jackson ImmunoResearch) and 4′,6-diamidino-2-phenylindole (DAPI) diluted in PBS with 0.1% Triton X-100 and 0.1% Tween-20 for 90 min at room temperature. The sections were then washed as described above and mounted with Aquamount mounting medium (18606, Polysciences) and imaged using Olympus FV1000 confocal microscope and quantified with the Imaris software.

### Whole mount immunostaining

For whole mount immunostaining, previously published protocol was followed with minor modifications^35^. Briefly, ~3 mm wide strips were cut from the periphery of cranial or left lobes. Strips were blocked in PBS with 0.3% Triton X-100 and 5% normal donkey serum and then incubated with primary antibodies diluted in PBS with 0.3% Triton X-100 at 4°C overnight in an 1.7 ml tube. The next day, the strips were washed with PBS+1% Triton X-100+1% Tween-20 (PBSTT) on a rocker at room temperature for 1 hr and this wash was repeated 3 times. Secondary antibodies were diluted in PBS with 0.3% Triton X-100 and added to the strips for an overnight incubation on a rocker at 4°C. The third day, strips were washed was PBSTT as described and then fixed with 2% PFA in PBS for 3 hr. Strips were washed 3 times with PBS and mounted on slides using Aquamount (18606, Polysciences) with the flat side of the strips facing the coverslip. Z-stacks of 20-30 um thickness at 1 um step size were taken using Olympus FV1000 confocal microscope and quantified with the Imaris software.

### Transmission Electron Microscopy

Samples were fixed with a solution containing 3% glutaraldehyde plus 2% paraformaldehyde in 0.1 M cacodylate buffer, pH 7.3, then washed in 0.1 M sodium cacodylate buffer, treated with 0.1% Millipore-filtered cacodylate buffered tannic acid, post fixed with 1% buffered osmium tetroxide, and stained en bloc with 1% Millipore-filtered uranyl acetate. The samples were dehydrated in increasing concentrations of ethanol, infiltrated, and embedded in LX-112 medium. The samples were polymerized in a 60°C oven for approximately 3 days. Ultrathin sections were cut in a Leica Ultracut microtome (Leica, Deerfield, IL), stained with uranyl acetate and lead citrate, and examined in a JEM 1010 transmission electron microscope (JEOL, USA, Inc., Peabody, MA) at an accelerating voltage of 80 kV. Digital images were obtained using AMT Imaging System (Advanced Microscopy Techniques Corp, Danvers, MA).

### Sendai virus infection

Viral infection was carried out following the previously described procedure^36^. Briefly, mice anesthetized with isoflurane were held by their upper incisors and subjected to oropharyngeal instillation of a non-lethal dose of Sendai Virus (ATCC #VR-105, RRID:SCR_001672CSCSSCS), approximately 2.1 × 10^7^ plaque-forming units, suspended in 40 µl of PBS. The control group received 40 µl of PBS.

### Tissue dissociation and fluorescence-activated cell sorting

Mouse lungs were collected as described above and subjected to our published protocol with minor modifications^37^. Briefly, connective tissues and trachea were removed, and lungs were minced to small pieces using forceps. The lungs were digested at 37°C for 30 min in 1.35 ml Liebovitz media (Gibco, 21083-027) with the following enzymes: 2 mg/ml collagenase type I (Worthington, CLS-1, LS004197), 0.5 mg/ml DNase I (Worthington, D, LS002007), and 2 mg/ml elastase (Worthingon, ESL, LS002294). To stop the enzymatic reaction, 300 ul fetal bovine serum (FBS, Invitrogen, 10082-139) were added to a final concentration of 20%. The digested tissue was mixed at least 10 times with pipette and filtered through a 70 µm cell strainer (Falcon, 352350) on ice in a 4°C cold room and transferred to a 2 ml tube. The cells were pelleted at 1537 rcf for 1 min. Then the supernatant was removed and 1 ml of red blood cell lysis buffer (15 mM NH_4_Cl, 12 mM NaHCO_3_, 0.1 mM EDTA, pH 8.0) was added and samples incubated on ice for 3 min. Cells were pelleted again by centrifugation at 1537 rcf for 1 min, washed with once Liebovitz + 10% FBS, resuspended with 1 ml Liebovitz + 10% FBS and filtered through 35 um cell strainer into a 5 ml glass tube. Samples for scMultiome were stained with CD45-PE/Cy7 (BioLegend, 103114), ECAD-PE (BioLegend, 147304), and ICAM2-A647 (Invitrogen, A15452) antibodies (1:250 dilutions for all antibodies) as well as SYTOX Blue (1:1000, Invitrogen, S34857) for 30 min on ice. Then samples were washed and resuspended with Liebovitz + 10% FBS, filtered again through a 35 um strainer into a 5 ml glass tube and sorted using Aria II Cell sorter with a 70 μm nozzle. Cell sorting data were analyzed using FlowJo 10.7. Lung epithelial cells from littermate *Cebpa*^F/F^; *Rosa^Sun1GFP/+^; Sftpc^CreER/+^*(n=2) and *Cebpa^F/+^; Rosa^Sun1GFP/+^; Sftpc^CreER/+^*lungs (n=2) were purified with a cell viability >77%. Single cell libraries were prepared using the Single Cell Multiome ATAC + Gene Expression kit (10x Genomics) and 10,000 nuclei were loaded per lane.

### Nuclei sorting for cell-type specific ChIP-seq

Lungs were collected as described above. Nuclei isolation followed the previously published protocol^8^. Briefly, lungs were minced with forceps and transferred to a 5 ml glass tube and crosslinked with 2 ml of 1% formaldehyde diluted with PBS from 10% buffered formalin (i.e. 3.7% formaldehyde;Thermo Fisher Scientific, 23-245-685) for 10 min at room temperature on a rocker. The excess formaldehyde was quenched by adding 0.5 M glycine (pH 5.0) to 0.125 mM final concentration and incubated for 15 min at RT on a rocker. The tissue was washed twice with ice cold PBS and resuspended and homogenized with Douncer homogenizer for 5 strokes in 1 ml of Isolation of Nuclei Tagged in specific Cell Types (INTACT) buffer (20 mM HEPES pH 7.4, 25 mM KCl, 0.5 mM MgCl_2_, 0.25 M sucrose, 1 mM DTT, 0.4% NP-40, 0.5 mM Spermine, 0.5 mM Spermidine)^33^ with protease inhibitor cocktail (cOmplete ULTRA Tablets, Mini, EDTA-free, EASY pack, Sigma, 5892791001). The tissue was then filtered through a 70 um strainer and then transferred to a 2 ml tube coated with 10 mg/ml BSA (Sigma, A3059). The nuclei were centrifuged at 384 rcf for 5 min 4°C, resuspended with 1ml ice cold PBS plus protease inhibitor cocktail and then filtered through a 35 um strainer into a 5 ml glass tube coated with 10 mg/ml BSA. SYTOX Blue was added with at a 1:1000 dilution. SYTOX+ and GFP+ nuclei were collected on Aria II Cell sorter with a 70 μm nozzle at 4°C into a 5 ml glass tube coated with 10 mg/ml BSA and containing 300ul of 10 mg/ml BSA with 5x protease inhibitor cocktail in PBS. *Sftpc^CreER/+;^ Rosa^Sun1GFP/+^* mice yielded 1–2 million GFP+ nuclei per adult lung.

### Chromatin immunoprecipitation

#### Lysis of the nuclei and shearing of the chromatin

AT2-specfic chromatin immunoprecipitation (ChIP) was performed on sorted nuclei following the previously published protocol with minor modification^8^. Briefly, sorted nuclei were split to aliquots of 1 million in 1.7 ml tubes and pelleted by centrifugation at 6708 rcf for 10 min at 4°C. The supernatant was discarded and pellets were resuspended in 100 ul of nuclei lysis buffer plus proteinase inhibitor and incubated on ice for 15 min. Samples were sonicated using Diagenode Bioruptor (B01060010) precooled to 4°C for 36 cycles of 30s ON/30s OFF to achieve a DNA fragment size of ~200-500 bp. Samples were then centrifuged at 13148 rcf for 10 min at 4°C and the supernatant was transferred to new 1.7 ml tubes. 20 ul of samples were saved as the input control in a separate tube and stored in −20°C till the reverse crosslinking step.

#### Bead Preparation

Two sets of Protein G Dynabeads (Thermo Fisher Scientific, 10004D) were washed twice with 1ml ChIP dilution buffer (16.7 mM Tris-HCl pH 8.1, 1.2 mM EDTA, 1.1% Triton X-100, 0.01% SDS with 1× protease inhibitor cocktail) and then blocked with 200 µl 20 mg/ml BSA (Jackson ImmunoResearch, 001-000-161), 4 µl 10 mg/ml salmon sperm DNA (Invitrogen, 15632-011) in the ChIP dilution buffer. The first set of Protein G Dynabeads was blocked for 1 hr on rotator at 4°C and was used to preclear the chromatin (40 ul of protein G Dynabeads per sample). The second set was blocked overnight (40 ul of protein G Dynabeads per sample) and was used for immunoprecipitation the second day.

#### Preclearing and antibody incubation

After chromatin shearing and centrifugation, samples were diluted to 1 ml with ChIP dilution buffer. The first set of beads were washed twice with ChIP dilution buffer using a magnetic adaptor before adding to samples, which were then incubated on rotator at 4°C for 1 hr. Using a magnetic adaptor, precleared samples were transferred to a new 1.7 tube and incubated with adequate amount of antibody overnight at 4°C on a rotator

#### Immunoprecipitation

The next day, the second set of beads were washed twice with ChIP dilution buffer and was added to the samples (chromatin/Ab solution) and incubated 3 hr on a rotator at 4°C. The sample were sequentially washed with 1 ml of the following prechilled buffers: low salt buffer (150 mM NaCl, 2 mM EDTA, 1% Triton X-100, 20 mM Tris-HCl pH 8.1, 0.1% SDS), high salt buffer (500 mM NaCl, 2 mM EDTA, 1% Triton X-100, 20 mM Tris-HCl pH 8.1, 0.1% SDS), lithium chloride buffer (250 mM LiCl, 1 mM EDTA, 1% NP-40, 10 mM Tris-HCl pH 8.1, 0.1% sodium deoxycholate), and TE buffer (10 mM pH 8.0 Tris and 1 mM EDTA) twice. Samples were resuspended in 300 ul TE buffer.

#### Reversal of crosslinking and DNA purification

The frozen inputs were thawed and diluted to 300 ul with TE buffer. Then samples and inputs were incubated for 4 hr at 37°C with 1.5 µl of 10 mg/ml RNase A (Qiagen, 1007885) and 15 µl 10% sodium dodecyl sulfate and 3.5 µl 20 mg/ml Proteinase K (Thermo Fisher, EO0491). Then samples were switched to 65°C overnight incubation. The next day, 300 ul of phenol: chloroform:isoamyl alcohol solution (Sigma, P2069-400ML) was added to samples and input and mixed by full-speed vortexing for ~2 s, then transferred to phase-lock tubes (Qiagen MaXtract, 129046) and centrifuged for 5 min at RT at 13148 rcf. 300 ul of chloroform was added to samples and mixed by inversion then centrifuged again at 13148 rcf for 5 min. The top liquid phase was transferred to a new 1.7 ml tube containing 2 µl of 20 µg/µl glycogen (Invitrogen, 10814-010). Then 600 µl of 100% ethanol and 30 µl of 3 M NaCl were added and mixed by quick vortexing, then samples were stored at −20°C overnight. Samples were centrifuged at 13148 rcf for 10 min at 4°C, then supernatant was discarded. The pellets were rinsed with 1 ml of 100% ethanol and centrifuged at 13148 rcf for 1 min at 4°C. The liquid was discarded, and pellets were air dried for 5 min and dissolved in 8 ul nuclease-free H_2_O.

#### ChIP-seq library preparation

DNA quantity was measured using a Qubit dsDNA HS Assay Kit (Invitrogen, Q23851). Then <5 ng ChIP sample DNA or <20 ng input DNA was used for sequencing libraries using the NEB Next Ultra II DNA Library Prep Kit for Illumina (New England BioLabs, E7645). In Step 1.3 (End Prep), thermocycler condition was modified as following: the heated lid set to ≥ 60°C, 30 min at 20°C then 60 min at 50°C; as decreasing the temperature to 50°C helps saving small fragments. The DNA was PCR amplified for 12 cycles using indexed primers (New England BioLabs, E7335S or E7500S) to barcode samples. Size selection and purification was achieved by double-sided (0.65 × −1× volume) using SPRIselect magnetic beads (Beckman Coulter, B23318). Concentrations were measured using the Qubit HS dsDNA assay. Samples were pooled with less than 20 barcoded samples per sequencing run on an Illumina NextSeq500.

### Bioinformatics analyses

#### ScRNA-seq time course analysis

Published scRNAseq dataset (GSE158192)^8^ compiled from 12-time points E14.5, E16.5, E18.5, P4, P6, P7, P8, P10, P15, P20, 10-week-old, and 15-week-old lung was analyzed using Seurat (v4.1). Cells were filtered out if they had a gene count of less than 200 or over 5000. Epithelial, immune, mesenchymal, and endothelial lineages were identified based on the expression of *Cdh1, Ptprc, Col3a1*, and *Icam2*, respectively. Doublets were filtered out and epithelial cells were subset and re-clustered. AT2 cells and SOX9 progenitors were subset from epithelial cells and used for downstream analysis. AT2-specific genes used in the heatmap were obtained by differential gene expression comparison between 15-wk AT2 and AT1 cells using Findmarkers resulting in 88 distinct genes. Subsequently, the expression level of these 88 genes was compared across different time points: E16.5, E18.5, and 15-wk. Genes were classified as early genes if they were present in at least 25% of E16.5 progenitor cells. The remaining genes were classified as late genes. Within this group, genes were classified as “mature” if the logFC between the 15-week time point and E18.5 exceeded 2 (i.e. 4-fold increase) (Table S1). Out of the 88 gene list, 6 genes were excluded (*Tpt1, Mt1, Mt2, Selenop, Wfdc2 and Ly6e*) as they met both early and mature criteria resulting in 82 genes that was used to generate heatmap in Fig. 1. Monocle 2.8.0 was used to analyze SOX9 progenitors and AT2 cells using the top 2000 genes to generate pseudo-time trajectories.

#### Pseudobulk ATAC-seq time course analysis

ScATAC-seq control datasets were from 5 different time points (E16.5, E18.5, P3, P8, and 9-wk). The P3 dataset was obtained from ^38^. Cell barcodes of AT2 or SOX9 progenitors were identified based on *Nkx2-1* and *Sftpc* or *Sox9* accessibility. Each sample was randomly divided cell barcodes into two to generate replicates. Then Sinto (version 0.4.0) was used to subset the *–F 2048 –q 30*. Peaks were called using MACS2 (v2.1.2) commands: *q 0.05 –nomodel –shift −100 –extsize 200 --broad*. Peaks overlapping with the mm10 blacklist were removed using bedtools. All overlapping peaks (>1kb) from different time points were merged using bedtools (v2.30) to obtain a reference peak set which was used for counting raw reads for each time point using Rsubread (version 2.8.2) and differential analysis was carried out by DEseq2 (version 1.34.0). Regions with (log2FC >1, FDR <0.01) were kept for downstream analysis. Diffbind (version 3.4.11) was used to obtain differentially accessible peaks representing the 4 categories: early lost (E16.5 vs E18.5); late lost (E18.5 vs 9-wk); early gain (E18.5 vs E16.5); and late gain (9-wk vs E18.5). GO analysis for each ATAC-seq cluster using Genomic Regions Enrichment of Annotation Tool (GREAT, version 4.0.4)^39^ using the default setting and the whole mouse genome (GRCm38/mm10) as the background.

#### Analysis of ChIP-seq data

Reads were concatenated and their quality was assessed by FastQC (v0.11.8) C (http://www.bioinformatics.babraham.ac.uk/ projects/fastqc/). Trimmomatic^40^ was used to filter poor-quality reads and trim poor-quality bases. Then reads were aligned to the mm10 reference genome using Bowtie2 (v.2.4.1) with the following parameters: -m1 -k1 -v1. Samtools (v1.15.) was used to convert aligned sam files to bam files. Then files were deduplicated and filtered for unmapped reads and low-quality alignments using Picard’s MarkDuplicates and samtools settings: *-b -h -F 4 -F 1024 -F 2048 -q 30*. Peaks were called using MACS2 with NarrowPeak setting *-g mm -n -B* for NKX2-1 and CEBPA. Peaks then were filtered for sites overlapping with the mm10 blacklist. Differential binding for NKX2-1 and CEBPA was carried out with Diffbind (v3.4.11) normalized for sample read depth and at a fixed peak width of 500 bp between controls and mutants. Bigwig files were generated using bamCoverage (version 3.3.2) with the following setting: *—binSize 20 —normalizeUsing BPM —smoothLength 60 —centerReads*. Heatmaps, genomic trackers and scatter maps were generated using EaSeq (version 1.2) (http://easeq.net)^41^. ChIP-Seq peaks were assigned to the nearest gene using ChIPseeker (version 1.3). CEBPA and NKX2-1 co-bound peaks were identified using ChIPpeakAnno_v3.2 using findOverlapsOfPeaks with max gap 50pb between peaks. Two replicates for each antibody were intersected using bedtools with the *-wa -wb* parameter.

### Motif analyses

De novo motif analysis was obtained using Homer (v. 4.10)^42^ ‘findMotifsGenome.pl -size 200 - mask’ with random backgrounds. Histograms of motif densities was performed using Homer ‘annotatePeaks’. To calculate the average distance between NKX and CEBP motifs, we used Homer ‘annotatePeaks.pl -size 200’ to search for NKX and CEBP motifs in NKX2-1/CEBPA co-bound peaks and calculated the distance from peak centers. ChromVAR (v1.16.0) was used to test overrepresented motifs in differentially accessible peaks by calculating motif scores on the cell level. ChromVar z-score deviation scores was calculated for the top 100 motifs curated from JASPAR (2020 Version) motif database.

### Single-cell multiome analysis

Control and mutant samples were aggregated via “cellranger-arc count” and cellranger-arc aggr (v2.0.0) using a custom mm10 reference genome that contains the *Sun1GFP* transcript. Downstream analysis was carried out using Seurat R package (v4.3) for RNA and Signac (v1.9) for ATAC and combined with the weighted nearest neighbor method. The signature score was calculated via the Seurat module score function using our previously published 119 progenitor genes, 100 AT2 genes and 100 AT1 genes^43^. 89 transitional genes (DATPS score) were derived from Choi et.al^18^. Psudobulk ATAC profiles for AT2 cells were generated using Sinto as described. Mutant gained and lost peaks were obtained using Diffbind. Overlapped peaks were obtained using bedtools intersect with parameters *-wa -wb*. Unique peaks were obtained by bedtools subtract with parameters −A. To generate RNA-seq/ATAC-seq scatter plots, differentially accessible peaks between mutant and control AT2 cells excluding proliferative cluster were linked to genes via Signac’s function LinkPeaks using the method described by SHARE-seq^44^. A custom script inspired by a prior study^45^ was used to combine gene expression and chromatin accessibility.

## Supporting information

Table S1. Raw data for Fig. 1, S1.

Table S2. Raw data for Fig. 2, S2.

Table S3. Raw data for Fig. 3.

Table S4. Raw data for Fig. 4.

Table S5. Raw data for Fig. 5.

Table S6. Raw data for Fig. 6.

## ACKNOWLEDGEMENTS

We thank Dr. Danielle Little for assistance with ChIP-seq and data analysis and Anne Lynch for assistance with Sendai virus infection. We thank Dr. Harold Chapman for providing the *Sftpc^CreER^* mice. The University of Texas MD Anderson Cancer Center DNA Analysis Facility, Flow Cytometry and Cellular Imaging Core Facility, and High Resolution Electron Microscopy Facility are supported by the Cancer Center Support Grant (P30CA016672). This work was supported by the University of Texas MD Anderson Cancer Center Retention Fund, and National Institutes of Health R01HL130129 and R01HL153511 (JC).

## AUTHOR CONTRIBUTIONS

DH and JC designed research; DH performed research and analyzed data; DH and JC wrote the paper; all authors read and approved the paper.

## COMPETING INTERESTS

The authors declare no competing interests.

## SUPPLEMENTARY FIGURES

**Fig. S1.**
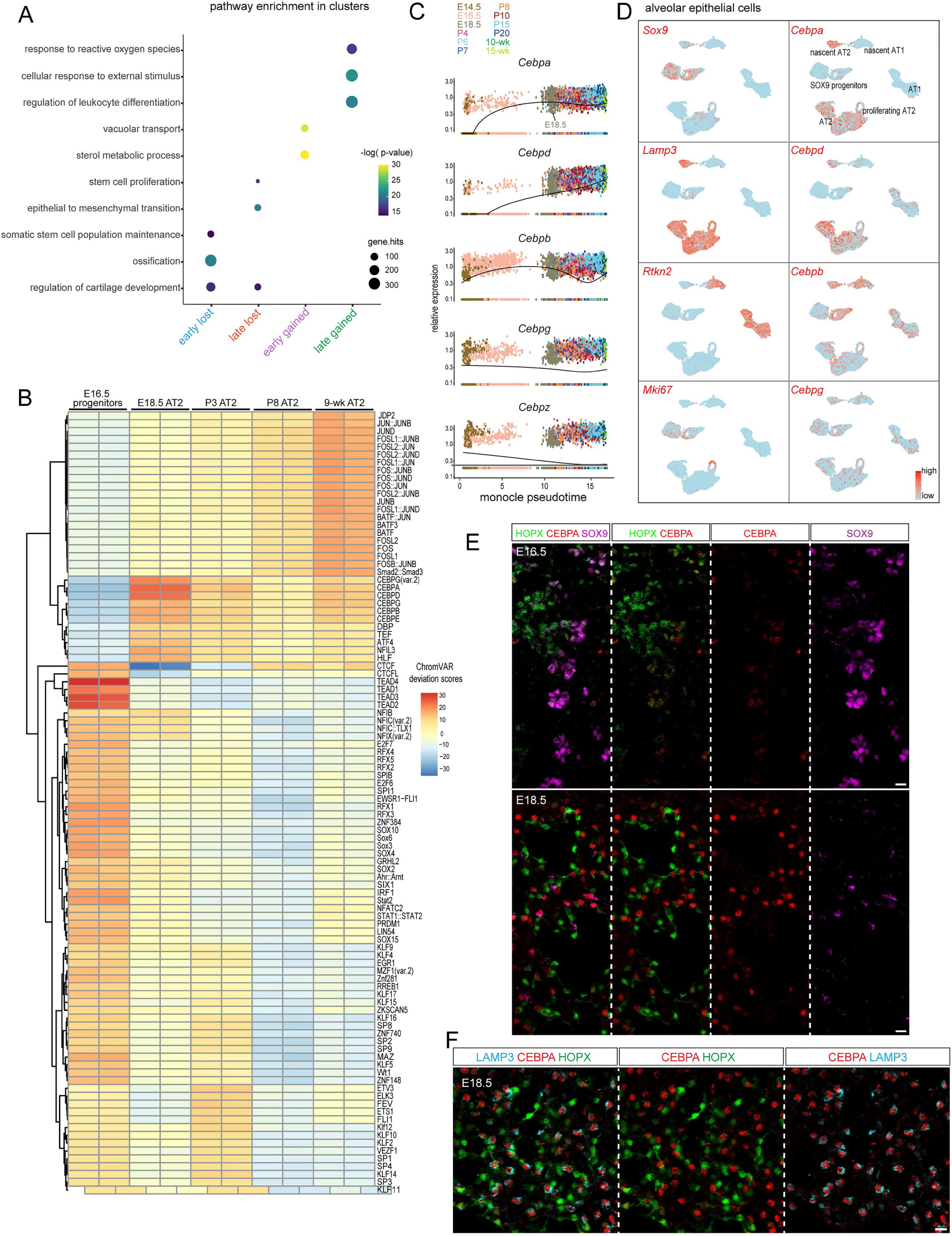
Time-course analysis of AT2 cell development and CEBPA expression. (**A**) Biological process GO terms for the nearest genes of the 4 ATAC-seq clusters in Fig. 1D. (**B**) Heatmap of ChromVAR deviation scores to the 100 most variable motifs across time. (**C**) Monocle pseudotemporal expression changes of 5 CEBP family members across 12 time points in Fig. 1B. *Cebpa*, but not other CEBP genes, reaches maximal expression upon AT2 specification. *Cebpe* is excluded due to lack of expression in alveolar epithelial cells. (**D**) Feature plots of Fig. 1A showing robust expression of *Cebpa*, but not other CEBP genes, in nascent AT2 cells. (**E**) Confocal images showing CEBPA is not expressed in SOX9 progenitors nor HOPX+ AT1 cells as SOX9 progenitors differentiate into AT1 and AT2 cells from E16.5 to E18.5. (**F**) Confocal images showing CEBPA is expressed in LAMP3+ AT2 cells but not HOPX+ AT1 cells. Scale: 10 um. See Table S1 for raw data.

**Fig. S2.**
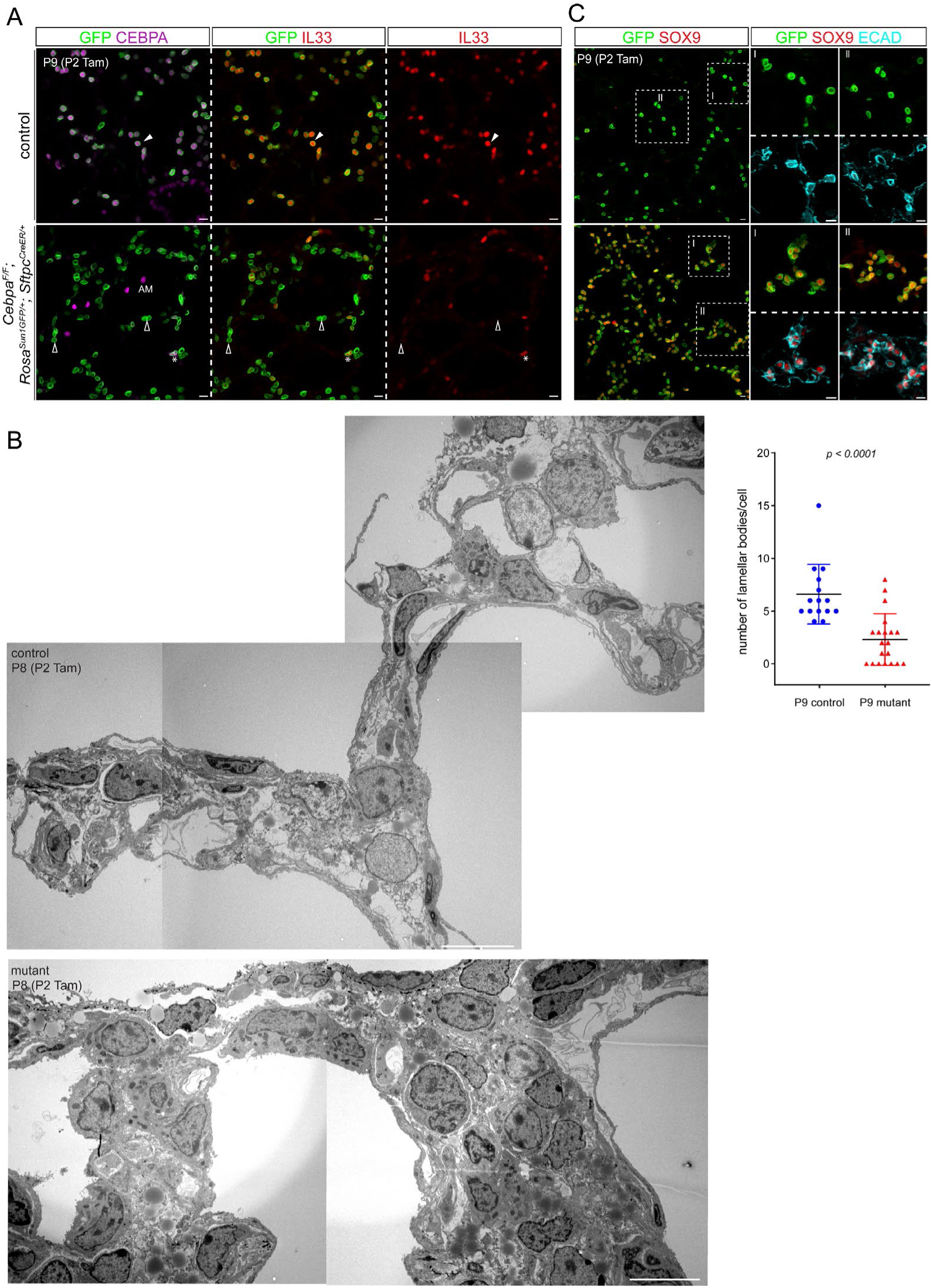
Characterization of neonatal *Cebpa* mutant AT2 cells. (**A**) Confocal images showing loss of CEBPA and IL33 in GFP+ recombined neonatal mutant AT2 cells (filled vs open arrowhead). AM, alveolar macrophage; *, escaper of *Cebpa* deletion still expressing IL33. (**B**) Stitched TEM images showing higher cell density in the mutant. Quantification of lamellar bodies for Fig. 2C (Student’s t-test). (**C**) Confocal images showing adjoining (ECAD) ectopic SOX9 cells in the mutant, resembling SOX9 progenitors at embryonic branch tips. Scale: 10 um. See Table S2 for raw data.

**Fig. S3.**
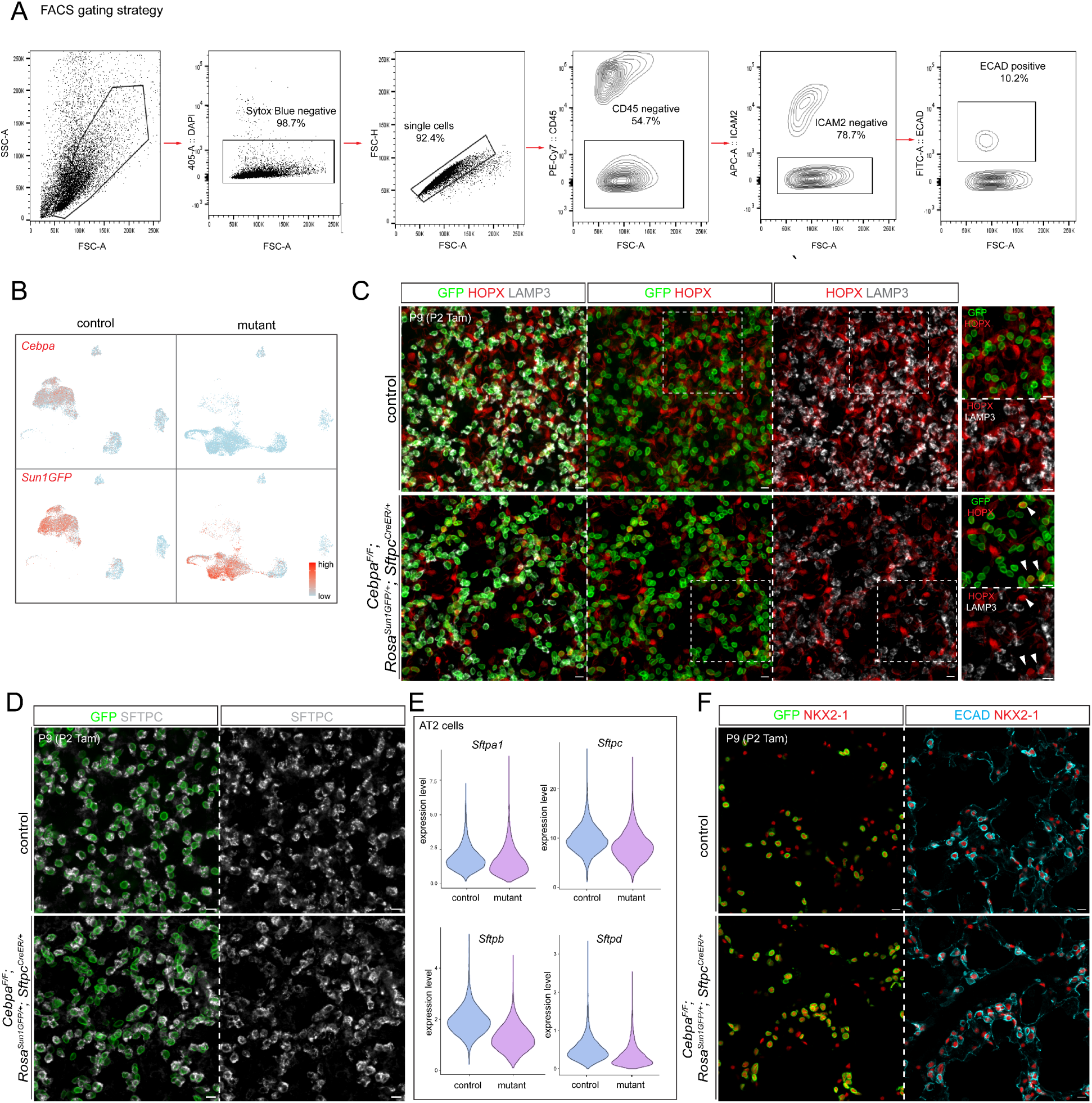
Multiome and staining of neonatal *Cebpa* mutant AT2 cells. (**A**) FACS gating strategy to purify lung epithelial cells. (**B**) Split feature plots of Fig. 3C to better visualize the control and mutant. (**C**) Confocal images showing that HOPX+ mutant AT2 cells do not express LAMP3 (arrowhead). (**D**) Confocal images showing persistent, albeit somewhat lower, SFTPC in mutant AT2 cells. (**E**) Violin plots of control and mutant AT2 cells in Fig. 3A showing a small decrease in surfactant gene expression. (**F**) Confocal images showing normal NKX2-1 expression in mutant AT2 cells. Scale: 10 um.

**Fig. S4.**
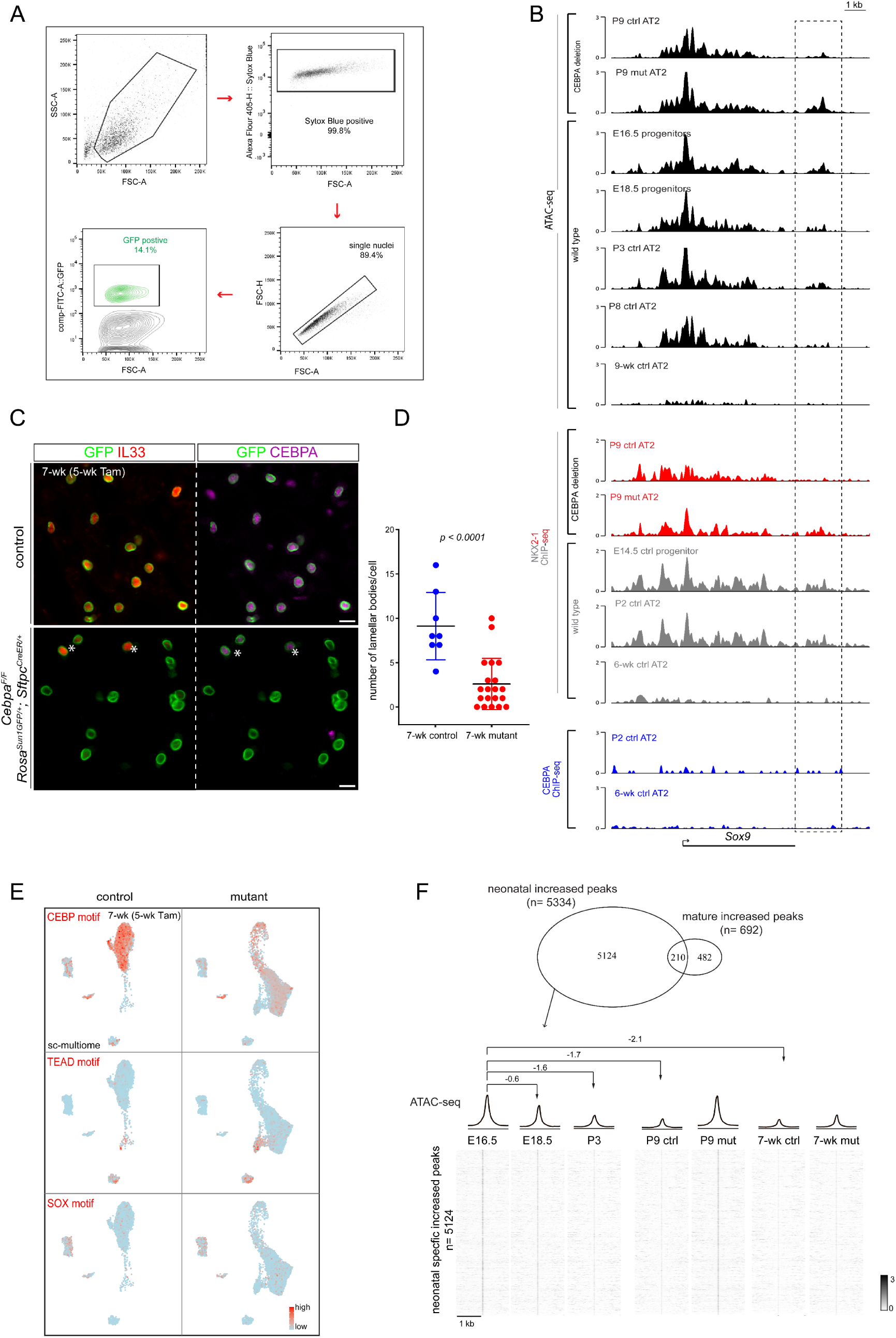
ChIP-seq and comparison of neonatal vs mature *Cebpa* mutant AT2 cells. (**A**) FACS gating strategy to purify AT2 nuclei for ChIP-seq. (**B**) Coverage plots showing a putative regulatory region 3’ to *Sox9* (box) that opens with more NKX2-1 binding upon *Cebpa* deletion, gradually closes and loses NKX2-1 binding during AT2 cell development in wild type lungs, and does not have CEBPA binding. (**C**) Confocal images showing loss of CEBPA and IL33 in GFP+ recombined mature mutant AT2 cells, except for escapers of deletion (asterisk). Scale: 10 um. (**D**) Quantification of lamellar bodies in mature AT2 cells for Fig. 5B (Student’s t-test). (**E**) Feature plots of motif activities for Fig. 5C. (**F**) Top: Venn diagram comparison of increased peaks in neonatal (Fig. 3F) vs mature (Fig. 5G) mutant AT2 cells. Bottom: heatmaps and profile plots showing that neonatal specific increased peaks gradually lose accessibility (log2 fold change) from E16.5 to 7-wk.

**Fig. S5.**
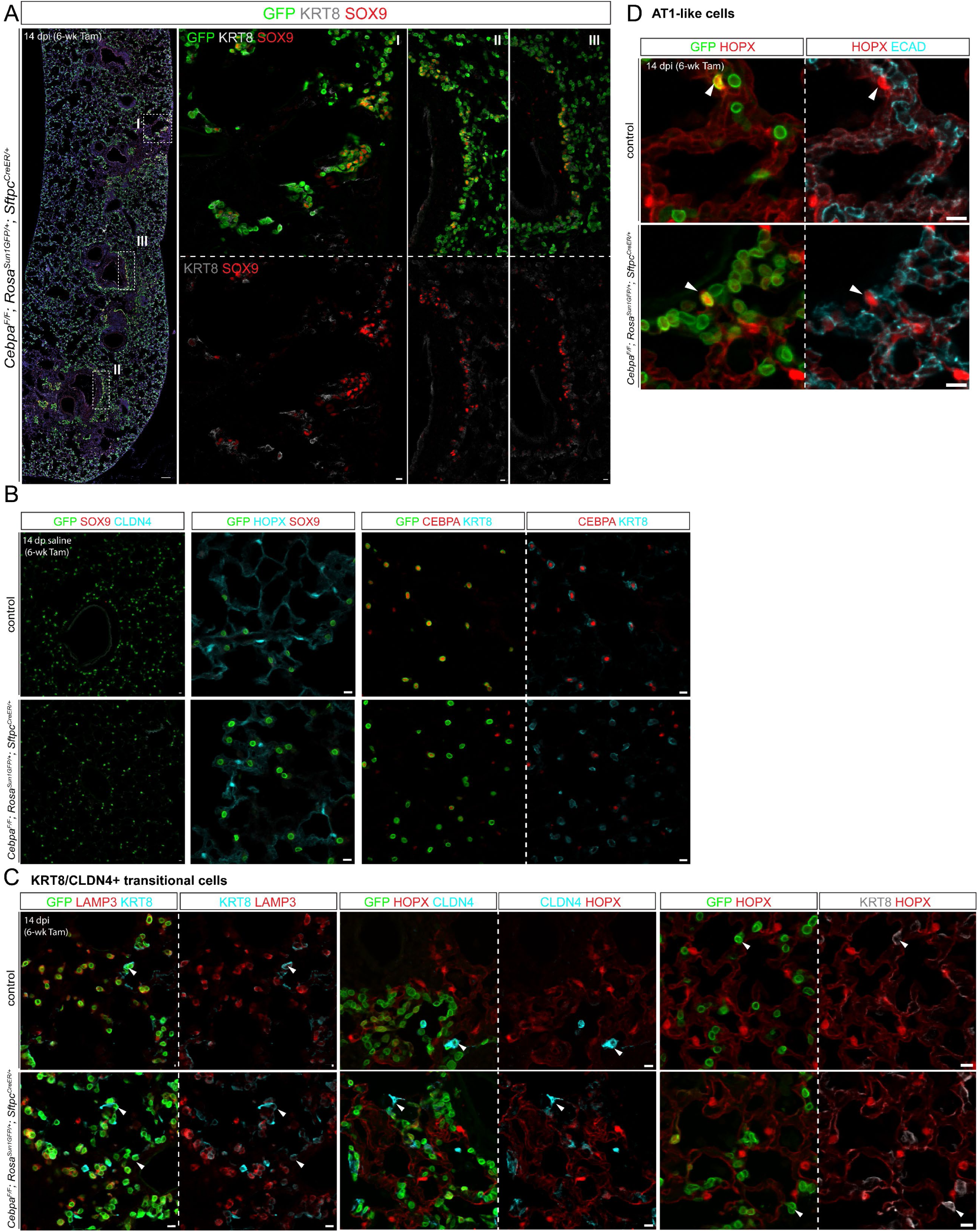
Characterization of control and *Cebpa* mutant lungs exposed to Sendai virus or saline. (**A**) Confocal images showing SOX9 reactivation, distinct from KRT8 expression, near lobe edges (I) and airways/macro-vessels (II, III). Scale: 100 um (10 um for insets). (**B**) Confocal images showing no SOX9 reactivation, HOPX expression, nor high KRT8 expression upon saline administration in control and mutant lungs. Baseline KRT8 expression is present in all AT2 cells. Scale: 10 um. (**C**) Confocal images showing that KRT8/CLDN4+ cells have low LAMP3 and no HOPX (arrowhead). Scale: 10 um. (**D**) Confocal images showing that AT1-like cells expressing HOPX (arrowhead) are no longer cuboidal (ECAD). Scale: 10 um.

**Fig. S6.**
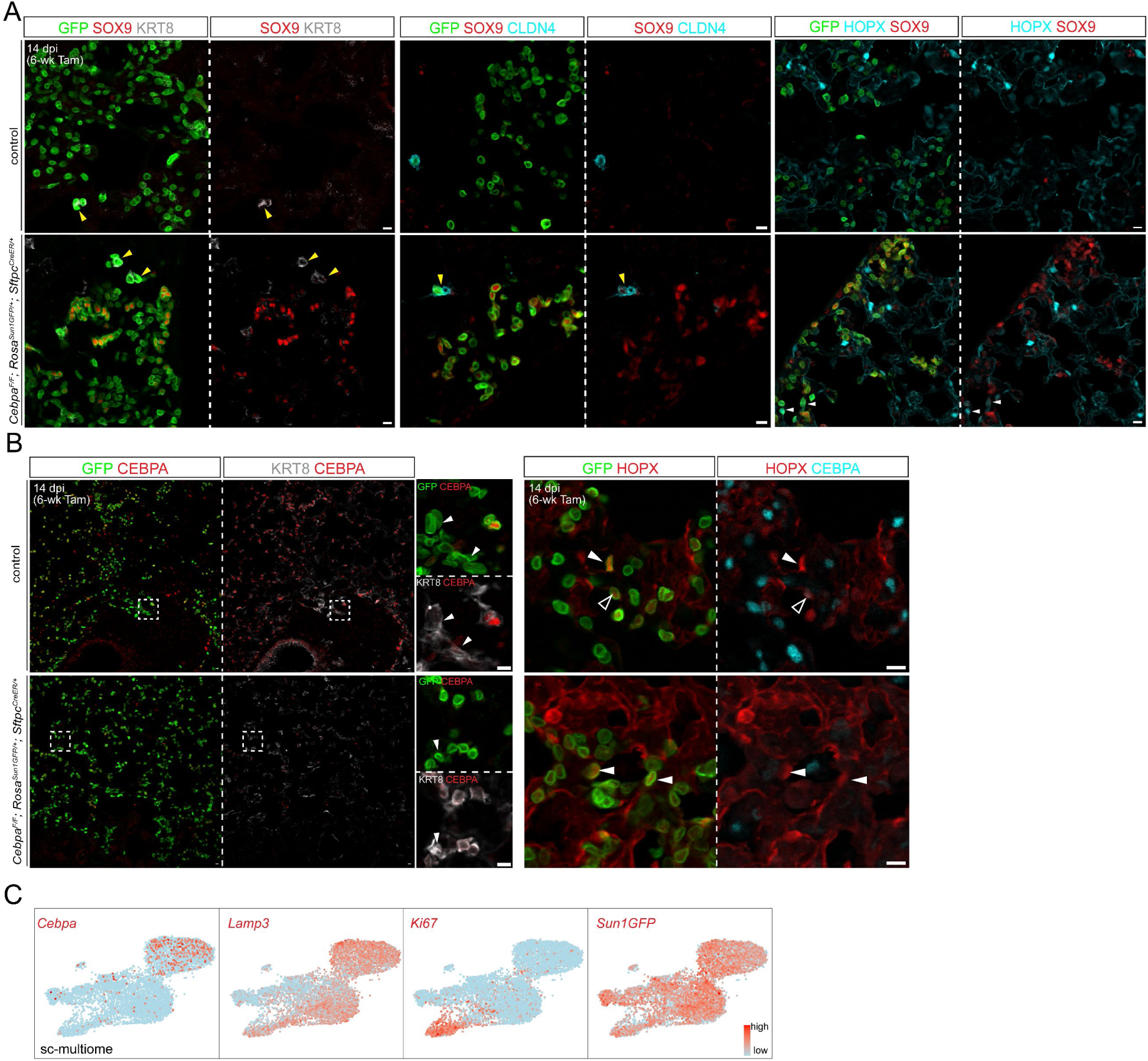
Additional characterization of SOX9 and CEBPA in infected control and *Cebpa* mutant lungs. (**A**) Confocal images showing that reactivated SOX9 in infected mutant AT2 cells is not in KRT8/CLDN4+ cells (yellow arrowhead) nor AT1-like cells (HOPX+; white arrowhead). Scale: 10 um. (**B**) Confocal images showing loss of CEBPA in KRT8/CLDN+ cells (left) and AT1-like cells (HOPX+; arrowhead) (right) even in the control lung. Open arrowhead, low CEBPA. Scale: 10 um. (**C**) Feature plots for Fig. 6G showing loss/reduction of *Cebpa* and *Lamp3* in KRT8/CLDN4+ and AT1-like cells even in the control lung.

## SUPPLEMENTARY TABLES

**Table S1. Raw data for Fig. 1, S1.**

**Table S2. Raw data for Fig. 2, S2.**

**Table S3. Raw data for Fig. 3.**

**Table S4. Raw data for Fig. 4.**

**Table S5. Raw data for Fig. 5.**

**Table S6. Raw data for Fig. 6.**

## Notes

### Competing Interest Statement

The authors have declared no competing interest.

## REFERENCES

1 Takahashi, K. & Yamanaka, S. Induction of pluripotent stem cells from mouse embryonic and adult fibroblast cultures by defined factors. Cell 126, 663–676, doi:10.1016/j.cell.2006.07.024 (2006).

2 Perez-Gonzalez, A., Bevant, K. & Blanpain, C. Cancer cell plasticity during tumor progression, metastasis and response to therapy. Nat Cancer 4, 1063–1082, doi:10.1038/s43018-023-00595-y (2023).

3 Ring, N. A. R., Valdivieso, K., Grillari, J., Redl, H. & Ogrodnik, M. The role of senescence in cellular plasticity: Lessons from regeneration and development and implications for age-related diseases. Dev Cell 57, 1083–1101, doi:10.1016/j.devcel.2022.04.005 (2022).

4 Lam, N. T. & Sadek, H. A. Neonatal Heart Regeneration: Comprehensive Literature Review. Circulation 138, 412–423, doi:10.1161/CIRCULATIONAHA.118.033648 (2018).

5 Desai, T. J., Brownfield, D. G. & Krasnow, M. A. Alveolar progenitor and stem cells in lung development, renewal and cancer. Nature, doi:10.1038/nature12930 nature12930 [pii] (2014).

6 Barkauskas, C. E. et al. Type 2 alveolar cells are stem cells in adult lung. J Clin Invest 123, 3025–3036, doi:10.1172/JCI68782 (2013).

7 Treutlein, B. et al. Reconstructing lineage hierarchies of the distal lung epithelium using single-cell RNA-seq. Nature, doi:10.1038/nature13173 (2014).

8 Little, D. R. et al. Differential chromatin binding of the lung lineage transcription factor NKX2-1 resolves opposing murine alveolar cell fates in vivo. Nat Commun 12, 2509, doi:10.1038/s41467-021-22817-6 (2021).

9 Helsel, A. R. et al. ID4 levels dictate the stem cell state in mouse spermatogonia. Development 144, 624–634, doi:10.1242/dev.146928 (2017).

10 Hock, H. et al. Tel/Etv6 is an essential and selective regulator of adult hematopoietic stem cell survival. Genes Dev 18, 2336–2341, doi:10.1101/gad.1239604 (2004).

11 Martis, P. C. et al. C/EBPalpha is required for lung maturation at birth. Development 133, 1155–1164, doi:dev.02273 [pii] 10.1242/dev.02273 (2006).

12 Gokey, J. J. et al. MEG3 is increased in idiopathic pulmonary fibrosis and regulates epithelial cell differentiation. JCI Insight 3, doi:10.1172/jci.insight.122490 (2018).

13 Zhao, Y., Zhu, Z., Shi, S., Wang, J. & Li, N. Long non-coding RNA MEG3 regulates migration and invasion of lung cancer stem cells via miR-650/SLC34A2 axis. Biomed Pharmacother 120, 109457, doi:10.1016/j.biopha.2019.109457 (2019).

14 Hernandez, B. J. et al. Intermediary Role of Lung Alveolar Type 1 Cells in Epithelial Repair Upon Sendai Virus Infection. Am J Respir Cell Mol Biol, doi:10.1165/rcmb.2021-0421OC (2022).

15 Strunz, M. et al. Alveolar regeneration through a Krt8+ transitional stem cell state that persists in human lung fibrosis. Nature communications 11, 3559, doi:10.1038/s41467-020-17358-3 (2020).

16 Kobayashi, Y. et al. Persistence of a regeneration-associated, transitional alveolar epithelial cell state in pulmonary fibrosis. Nat Cell Biol 22, 934–946, doi:10.1038/s41556-020-0542-8 (2020).

17 Jiang, P. et al. Ineffectual Type 2-to-Type 1 Alveolar Epithelial Cell Differentiation in Idiopathic Pulmonary Fibrosis: Persistence of the KRT8(hi) Transitional State. Am J Respir Crit Care Med 201, 1443–1447, doi:10.1164/rccm.201909-1726LE (2020).

18 Choi, J. et al. Inflammatory Signals Induce AT2 Cell-Derived Damage-Associated Transient Progenitors that Mediate Alveolar Regeneration. Cell Stem Cell 27, 366–382 e367, doi:10.1016/j.stem.2020.06.020 (2020).

19 Little, D. R. et al. Transcriptional control of lung alveolar type 1 cell development and maintenance by NK homeobox 2-1. Proc Natl Acad Sci U S A 116, 20545–20555, doi:10.1073/pnas.1906663116 (2019).

20 Penkala, I. J. et al. Age-dependent alveolar epithelial plasticity orchestrates lung homeostasis and regeneration. Cell Stem Cell, doi:10.1016/j.stem.2021.04.026 (2021).

21 Shiraishi, K. et al. Biophysical forces mediated by respiration maintain lung alveolar epithelial cell fate. Cell 186, 1478–1492 e1415, doi:10.1016/j.cell.2023.02.010 (2023).

22 Gokey, J. J. et al. YAP regulates alveolar epithelial cell differentiation and AGER via NFIB/KLF5/NKX2-1. iScience 24, 102967, doi:10.1016/j.isci.2021.102967 (2021).

23 Srivastava, D. & DeWitt, N. In Vivo Cellular Reprogramming: The Next Generation. Cell 166, 1386–1396, doi:10.1016/j.cell.2016.08.055 (2016).

24 Sugimoto, K., Gordon, S. P. & Meyerowitz, E. M. Regeneration in plants and animals: dedifferentiation, transdifferentiation, or just differentiation? Trends in cell biology 21, 212–218, doi:10.1016/j.tcb.2010.12.004 (2011).

25 Gola, A. & Fuchs, E. Environmental control of lineage plasticity and stem cell memory. Curr Opin Cell Biol 69, 88–95, doi:10.1016/j.ceb.2020.12.015 (2021).

26 Blanco, M. A. et al. Chromatin-state barriers enforce an irreversible mammalian cell fate decision. Cell Rep 37, 109967, doi:10.1016/j.celrep.2021.109967 (2021).

27 Ito, K. & Zaret, K. S. Maintaining Transcriptional Specificity Through Mitosis. Annu Rev Genomics Hum Genet 23, 53–71, doi:10.1146/annurev-genom-121321-094603 (2022).

28 Macrae, T. A., Fothergill-Robinson, J. & Ramalho-Santos, M. Regulation, functions and transmission of bivalent chromatin during mammalian development. Nat Rev Mol Cell Biol 24, 6–26, doi:10.1038/s41580-022-00518-2 (2023).

29 Huyghe, A., Trajkova, A. & Lavial, F. Cellular plasticity in reprogramming, rejuvenation and tumorigenesis: a pioneer TF perspective. Trends in cell biology, doi:10.1016/j.tcb.2023.07.013 (2023).

30 Snyder, E. L. et al. Nkx2-1 represses a latent gastric differentiation program in lung adenocarcinoma. Mol Cell 50, 185–199, doi:10.1016/j.molcel.2013.02.018 S1097-2765(13)00174-3 [pii] (2013).

31 Zhang, P. et al. Enhancement of hematopoietic stem cell repopulating capacity and self-renewal in the absence of the transcription factor C/EBPα. Immunity 21, 853–863 (2004).

32 Chapman, H. A. et al. Integrin α6β4 identifies an adult distal lung epithelial population with regenerative potential in mice. The Journal of clinical investigation 121, 2855–2862 (2011).

33 Mo, A. et al. Epigenomic signatures of neuronal diversity in the mammalian brain. Neuron 86, 1369–1384 (2015).

34 Chen, J. & Nathans, J. Estrogen-related receptor β/NR3B2 controls epithelial cell fate and endolymph production by the stria vascularis. Developmental cell 13, 325–337 (2007).

35 Yang, J. et al. The development and plasticity of alveolar type 1 cells. Development 143, 54–65 (2016).

36 Hernandez, B. J. et al. Intermediary Role of Lung Alveolar Type 1 Cells in Epithelial Repair upon Sendai Virus Infection. Am J Respir Cell Mol Biol 67, 389–401, doi:10.1165/rcmb.2021-0421oc (2022).

37 Little, D. R. et al. Transcriptional control of lung alveolar type 1 cell development and maintenance by NK homeobox 2-1. Proceedings of the National Academy of Sciences 116, 20545–20555 (2019).

38 Penkala, I. J. et al. Age-dependent alveolar epithelial plasticity orchestrates lung homeostasis and regeneration. Cell Stem Cell 28, 1775–1789 e1775, doi:10.1016/j.stem.2021.04.026 (2021).

39 McLean, C. Y. et al. GREAT improves functional interpretation of cis-regulatory regions. Nature biotechnology 28, 495–501 (2010).

40 Bolger, A. M., Lohse, M. & Usadel, B. Trimmomatic: a flexible trimmer for Illumina sequence data. Bioinformatics 30, 2114–2120 (2014).

41 Lerdrup, M., Johansen, J. V., Agrawal-Singh, S. & Hansen, K. An interactive environment for agile analysis and visualization of ChIP-sequencing data. Nature structural & molecular biology 23, 349–357 (2016).

42 Heinz, S. et al. Simple Combinations of Lineage-Determining Transcription Factors Prime cis-Regulatory Elements Required for Macrophage and B Cell Identities. Molecular Cell 38, 576–589, doi:10.1016/j.molcel.2010.05.004 (2010).

43 Gerner-Mauro, K. N., Akiyama, H. & Chen, J. Redundant and additive functions of the four Lef/Tcf transcription factors in lung epithelial progenitors. Proc Natl Acad Sci U S A 117, 12182–12191, doi:10.1073/pnas.2002082117 (2020).

44 Ma, S. et al. Chromatin potential identified by shared single-cell profiling of RNA and chromatin. Cell 183, 1103–1116.e1120 (2020).

45 Kiani, K., Sanford, E. M., Goyal, Y. & Raj, A. Changes in chromatin accessibility are not concordant with transcriptional changes for single-factor perturbations. Molecular Systems Biology 18, e10979 (2022).

